# Identifying loci under selection via explicit demographic models

**DOI:** 10.1101/2020.07.20.211581

**Authors:** Hirzi Luqman, Alex Widmer, Simone Fior, Daniel Wegmann

## Abstract

Adaptive genetic variation is a function of both selective and neutral forces. To accurately identify adaptive loci, it is thus critical to account for demographic history. Theory suggests that signatures of selection can be inferred using the coalescent, following the premise that genealogies of selected loci deviate from neutral expectations. Here, we build on this theory to develop an analytical framework to identify Loci under Selection via explicit Demographic models (LSD). Under this framework, signatures of selection are inferred through deviations in demographic parameters, rather than through summary statistics directly, and demographic history is accounted for explicitly. Leveraging on the property of demographic models to incorporate directionality, we show that LSD can provide information on the environment in which selection acts on a population. This can prove useful in elucidating the selective processes underlying local adaptation, by characterising genetic trade-offs and extending the concepts of antagonistic pleiotropy and conditional neutrality from ecological theory to practical application in genomic data. We implement LSD via Approximate Bayesian Computation and demonstrate, via simulations, that LSD has i) high power to identify selected loci across a large range of demographic-selection regimes, ii) outperforms commonly applied genome-scan methods under complex demographies, and iii) accurately infers the directionality of selection for identified candidates. Using the same simulations, we further characterise the behaviour of isolation-with-migration models conducive to the study of local adaptation under regimes of selection. Finally, we demonstrate an application of LSD by detecting loci and characterising genetic trade-offs underlying flower colour in *Antirrhinum majus*.

## 1 INTRODUCTION

Elucidating the genetic basis of adaptation and identifying genetic determinants of population and species divergence are key foci in evolutionary biology. In natural systems, genetic variation is shaped by the demographic history (driven by the neutral processes of mutation, migration and drift) together with natural selection on loci underlying adaptive traits. While all gene genealogies are constrained by the demographic history of the population, the genealogies of loci affected by selection are perturbed and may differ in key characteristics compared to those evolving under neutrality, though converging patterns can arise (Bierne, Welch, Loire, Bonhomme, & David, 2011; Edmonds, Lillie, & Cavalli-Sforza, 2004; Laurent Excoffier, Foll, & Petit, 2009; J. Li et al., 2012; Montgomery Slatkin & Excoffier, 2012). Disentangling the genomic signatures generated by these two processes, i.e. correctly identifying adaptive loci, remains a prevailing challenge in the field of population genetics (Biswas & Akey, 2006; Horscroft, Ennis, Pengelly, Sluckin, & Collins, 2019; Luikart, England, Tallmon, Jordan, & Taberlet, 2003).

A multitude of methods have been developed that identify loci under selection as those whose summary statistics deviate from the genome-wide distribution. These “outlier” approaches can generally be grouped into three classes: those that 1) detect regions of elevated differentiation between populations (via e.g. *F*_ST_-related statistics), 2) detect regions of perturbed site frequency spectrum (SFS) via diversity or diversity-related estimators (e.g. π, Tajima’s *D*) and 3) detect regions of extensive linkage disequilibrium (LD) via haplotype statistics (e.g. EHH, iHS) (M. A. Beaumont & Nichols, 1996; Biswas & Akey, 2006; Luikart et al., 2003; Oleksyk, Smith, & O’Brien, 2010; Sabeti et al., 2002; Vitti, Grossman, & Sabeti, 2013). While in empirical studies inference of selection is often achieved through corroboratory evidence from multiple measures, the choice of a particular class and hence summary statistic is motivated by the type of selection one aims to infer; with the first geared towards divergent selection between populations and the others towards footprints of selection within single populations.

Under the premise that adaptive genetic variation is a function of both selective and neutral forces, accounting for the demographic history of the study system is critical for the correct identification of selected loci (L. Excoffier, Hofer, & Foll, 2009; François, Martins, Caye, & Schoville, 2016; Hoban et al., 2016; Hofer, Ray, Wegmann, & Excoffier, 2009). This is commonly achieved by contrasting locus-specific statistics against an estimate of the expected distribution of these statistics under demography alone, with the power of such an approach being a function of both the summary statistics used and the accuracy with which the neutral distribution is inferred. In the context of identifying local adaptation, the canonical statistics employed is *F_ST_*, and the first methods to infer its neutral distribution used simulations under an island model calibrated by matching the observed heterozygosity at each locus (M. A. Beaumont & Nichols, 1996; L. Excoffier et al., 2009). Under island models, the distribution of sample allele frequencies is also well captured by Pólya distributions (Balding & Nichols, 1994; Rannala & Hartigan, 1996), which can be learned using likelihood-based methods that jointly classify loci into neutral and selected classes (Mark A. Beaumont & Balding, 2004; Foll & Gaggiotti, 2008; Galimberti et al., 2020). While generally powerful, these methods suffer from high false-positive rates in the case of asymmetric divergence between populations (Galimberti et al., 2020; Lotterhos & Whitlock, 2014; Luu, Bazin, & Blum, 2017), which violates a key assumption of island models. This can be alleviated by using hierarchical island models (Foll, Gaggiotti, Daub, Vatsiou, & Excoffier, 2014; Galimberti et al., 2020) or a more flexible distribution to capture neutral allele frequencies (e.g. PCA, Luu, Bazin, & Blum, 2017). For some natural systems however, these approaches may still be insufficient to capture the demographic history of the population and a (potentially complex) demographic model should be used. Williamson et al. (2005), for instance, inferred such a model from putatively neutral loci and then identified loci under selection as those for which an additional selection parameter is required.

Rather than modelling selection explicitly, loci under selection may also be identified under pure demographic models through locus-specific demographic parameters, under the premise that the demographic parameters of selected loci are expected to deviate from neutral expectations (Barton & Bengtsson, 1986; Charlesworth, 2009; Charlesworth, Nordborg, & Charlesworth, 1997; Fusco & Uyenoyama, 2011; Galtier, Depaulis, & Barton, 2000; Gossmann, Woolfit, & Eyre-Walker, 2011; Petry, 1983; Sousa, Carneiro, Ferrand, & Hey, 2013). Under coalescent theory, demographic models are parametrized by the effective sizes (*N_E_*) of each population and the effective rates of migration (*M_E_*) between them, which respectively describe the level of drift and gene flow within and between populations (Charlesworth, 2009; Petry, 1983). Importantly, both *N_E_* and *M_E_* may change through time.

Different modes of selection and adaptive processes can be expected to alter these demographic parameters in different ways. In a single population, a selective sweep is expected to reduce *N_E_* at selected and linked sites while diversifying selection is expected to increase it (Galtier et al., 2000; Gossmann et al., 2011). In the case of two or more populations connected by gene flow, balancing selection and adaptive introgression are expected to increase *M_E_* at selected and linked sites, while divergent selection is expected to reduce *M_E_* at those sites (Charlesworth et al., 1997; Geraldes, Ferrand, & Nachman, 2006; Petry, 1983; Won, Sivasundar, Wang, & Hey, 2005).

Notably, locus-specific demographic parameters are not just informative about the strength (i.e. the magnitude of variation in *N_E_* or *M_E_*) and mode of selection (i.e. a reduction or elevation of *N_E_* or *M_E_*), but may also identify the population (or environment) in which selection acts (i.e. a reduction in *M_E_* in one but not the other direction). Directional selection modulates fitness of natural populations by purging maladaptive alleles via extrinsic barriers such as hybrid or immigrant inviability, or lower fecundity (Naisbit, Jiggins, & Mallet, 2001; Nosil, Vines, & Funk, 2005; Rundle & Whitlock, 2001; Schluter, 2000). This effectively reduces *M_E_* at selected loci proportionally to the strength of selection (Petry, 1983). Under local adaptation, alternate alleles may confer higher fitness in their respective local environment but reduced fitness in the foreign environment, i.e. antagonistic pleiotropy (AP), or an allele may confer higher fitness in its local environment but have no differential effect relative to the alternative allele in the foreign environment, i.e. conditional neutrality (CN) (Anderson, Lee, Rushworth, Colautti, & Mitchell-Olds, 2013; Kawecki & Ebert, 2004; Savolainen, Lascoux, & Merilä, 2013). To date, such genetic trade-offs have only been characterized in a handful of cases, primarily through demanding experiments involving the transplanting of alternate genotypes in their reciprocal environments (Anderson, Lee, & Mitchell-Olds, 2014; Anderson et al., 2013; Oakley, Ågren, Atchison, & Schemske, 2014; Troth, Puzey, Kim, Willis, & Kelly, 2018). Characterizing the nature of such trade-offs directly from genomic data presents a promising complementary approach, applicable to natural populations, to investigate the genetic basis of local adaptation, a key concept in ecological genetics.

Inferring demographic parameters using coalescent theory is, however, computationally challenging, as the underlying but unknown genealogies must be integrated out numerically (Hey & Nielsen, 2007). As a result, there exists only a single likelihood implementation to infer locus-specific and global demographic parameters jointly: an MCMC sampler that attributes loci to different classes (e.g. selected and neutral) and jointly infers the demographic parameters of a two-population isolation-with-migration (IM) model for each group (Sousa et al., 2013). To extend this approach to more complex models, simulation-based techniques such as Approximate Bayesian Computation (Mark A. Beaumont, Zhang, & Balding, 2002; Marjoram & Tavaré, 2006; Sisson, Fan, & Beaumont, 2018) may be employed. The use of ABC to infer genome-wide demographic parameters has a long tradition (e.g. Dussex, Wegmann, & Robertson, 2014; Wegmann & Excoffier, 2010) and it may also be used to infer locus-specific parameters in a hierarchical setting, but it is computationally challenging. Indeed, the dimensionality of a genome-wide model is prohibitively large for any naïve Monte-Carlo scheme as the probability that a simulation matches the data at all loci is virtually zero. When inferring genome-wide parameters, loci are exchangeable and this problem is easily overcome by using summary statistics that are functions of all loci such as the (scaled) moments of the distribution of locus-specific statistics (Tavaré, Balding, Griffiths, & Donnelly, 1997; Wegmann, Leuenberger, & Excoffier, 2009). To benefit from this in a hierarchical setting, Bazin *et al*. (2010) proposed a two-step algorithm in which hierarchical parameters are inferred based on moments of locus-specific summary statistics, and locus-specific parameters are then inferred with simulations of a single locus conducted with parameters drawn from the posterior distribution of the hierarchical parameters. As recently shown (Kousathanas, Leuenberger, Helfer, Quinodoz, & Foll, 2016), this approach can be generalized to arbitrary parameter dependencies when using an ABC-MCMC setting that also eliminates the need for a two-step approach. These approaches were successfully used to infer locus-specific (Bazin, Dawson, & Beaumont, 2010) and cluster-specific (Aeschbacher, Futschik, & Beaumont, 2013) migration rates, as well as locus-specific selection coefficients (Foll, Poh, et al., 2014; Kousathanas et al., 2016) for up to several hundred loci. Scaling such inference up to whole-genomes, however, remains difficult due to the requirement to simulate all loci.

In this paper, we introduce LSD, an alternative framework for identifying Loci under Selection via explicit Demographic models that scales to genomic data. Similar to the approach by Bazin *et al*. (2010), LSD works in two steps. But rather than inferring hierarchical parameters for all loci, LSD first obtains a point estimate of demographic parameters for neutral loci, which is then compared against per-locus estimates to identify selected loci. This has the benefit of requiring only simulations of a single locus, which can be efficiently recycled (Thalmann et al., 2011). As a downside, the approach requires *a priori* knowledge on putative neutral sites. However, as we show with simulations, the approach is very robust to mis-specifications.

While LSD is flexible regarding the choice of demographic model and can in principle accommodate any discrete population model (including single population and stepping-stone models) as well as detect different modes of selection, we demonstrate here LSD’s utility in studies of local adaptation by focusing on the detection of loci under divergent selection between populations under isolation-with-migration (IM) models. We validate and assess the performance of LSD via extensive simulations, provide general insights into the properties of IM models in relation to the power of LSD and other widely applied genome scan methods, and demonstrate an application of the method to the detection of functionally validated loci underlying flower colour in two parapatric subspecies of *Antirrhinum majus* (common snapdragon) (Schwinn et al., 2006; Tavares et al., 2018).

## 2 MATERIALS AND METHODS

### 2.1 Model

We begin by outlining the conceptual framework underlying LSD. Consider a demographic model 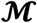, parameterized by demographic parameters ***θ***, that generates genetic data ***D***. To quantify deviations from neutrality, LSD first estimates the demographic parameters 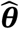 from a collection of loci assumed to be neutral (Figure S1). In a second step, LSD performs demographic inference on all loci and determines the posterior distribution *π_l_*(***θ***) = *π*(***θ|D_l_***) for each locus. Finally, LSD assesses the concordance of 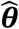 with *π_l_*(***θ***) by determining *h_l_*, the highest posterior density interval (HPDI) of ***π_l_***(***θ***) that contains 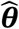, and uses *p_l_* = 1 – *h_l_* as a metric to identify locus *l* as incompatible with 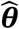. For loci with parameters 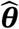, *p_l_* follows a uniform distribution (the coverage property) and is interpreted as a *p*-value to reject 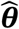 for locus *l*. The joint posterior distribution *π_l_*(***θ***) may further provide information on the magnitude and directionality of selection.

Given that the evaluation of the likelihood is non-trivial and may be intractable under more complex models, we resort to an approximate approach (Marjoram & Tavaré, 2006) (Figure 1). Under an Approximate Bayesian Computation (ABC) framework, the likelihood is approximated by simulations, the outcomes of which are compared with observed data in terms of summary statistics. That is, we find the set of parameters ***θ*** that minimise the distance between the observed data ***D*** and the simulated data ***D′***. To efficiently evaluate this, we reduce the dimensionality of the data via summarising them into a set of lower-dimensional summary statistics ***S*** and ***S′***, which are selected to capture the relevant information in ***D*** and ***D′***, respectively (Mark A. Beaumont et al., 2002; Joyce & Marjoram, 2008; Sisson et al., 2018).

**Figure 1.**
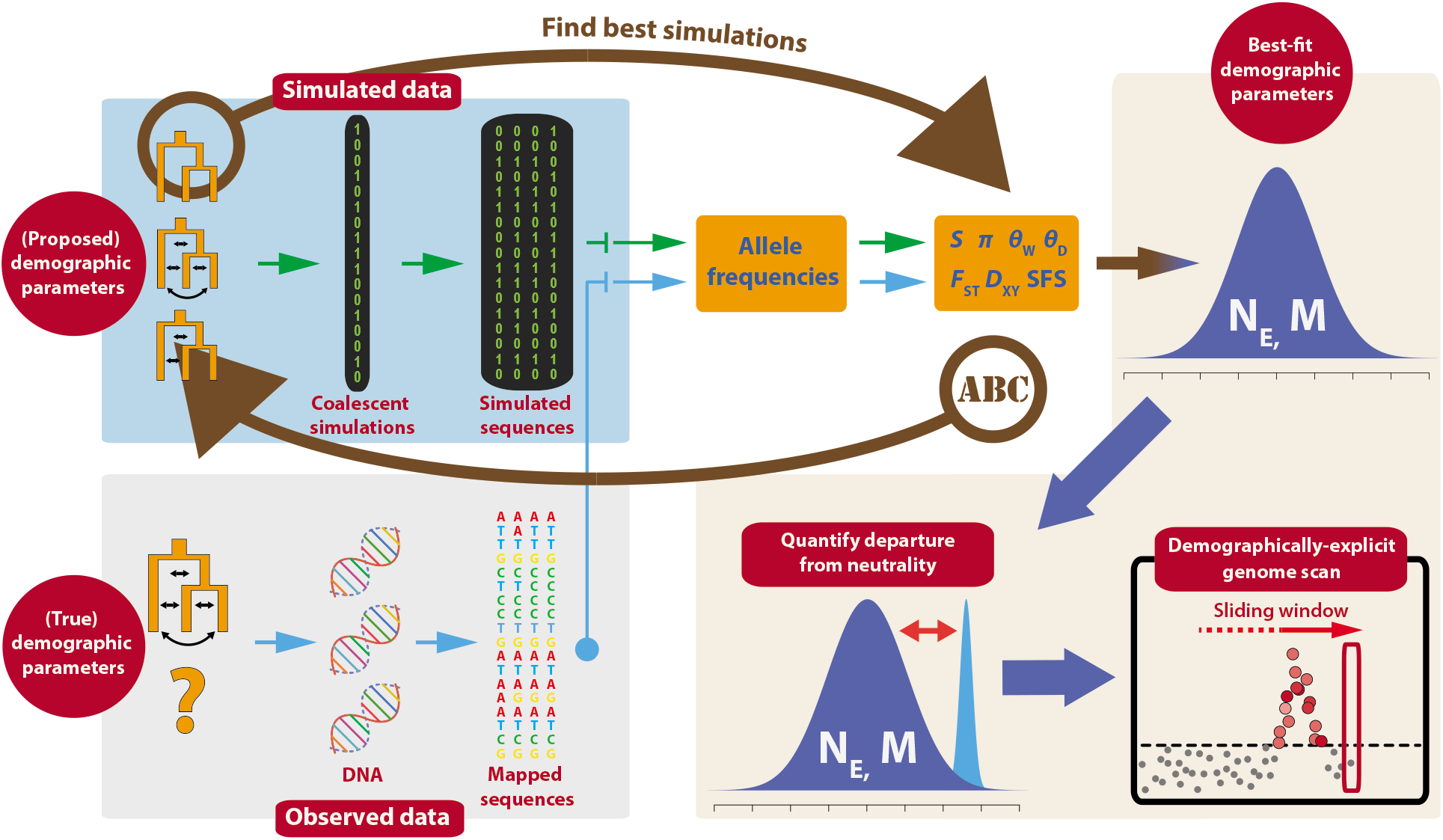
Analytical framework to identify Loci under Selection via explicit Demographic models (LSD). LSD identifies loci under selection by first estimating demographic parameters and then quantifying the departure of these parameters from neutral expectations. Our specific implementation of LSD employs Approximate Bayesian Computation (ABC) for parameter estimation, and is performed in a genome scan approach.

An appropriate model for generating simulated genetic data is provided by coalescent theory (Kingman, 1982; Wakeley, 2001), parametrised by population demographic parameters ***θ*** = {***N_E_, M_E_***, *μ*}, where ***N_E_*** refers to the vector of effective population sizes, ***M_E_*** to the vector of effective migration rates, and *μ* to the mutation rate. We stress that population sizes and migration rates may vary through time.

### 2.2 Implementation

We implemented the framework described above as shown in Figure 1 and detailed below.

#### Simulations

While the framework is readily used for any type of locus, we consider here a locus to consist of a genomic window with a shared genealogy. This effectively implies that recombination is free between loci and fixed within. We simulate genealogies using msms (Ewing & Hermisson, 2010), under a user-defined demographic model. The processing, format and final output of observed genetic data will often differ from that of coalescent simulations, given that observed genetic data may be subject to various pre-sequencing (e.g. pooling), sequencing (e.g. sequencing errors, stochastic sampling of reads) and post-sequencing (e.g. filters) events that perturb and reformat the data from the original source. We thus implement two complementary programs that interface with coalescent simulators to replicate observed sequencing pipelines and generate simulated sequencing data: *LSD-High* can accommodate and simulate both individual and pooled data and assumes mid to high coverage (>10x) data, while *LSD-Low* accepts individual data and can additionally accommodate low coverage (>2x) data by utilising genotype likelihoods via msToGLF and ANGSD (Korneliussen, Albrechtsen, & Nielsen, 2014). A suite of summary statistics is then calculated for the simulated and observed data via the same programs. Summary statistics currently implemented include the number of segregating sites (*S*), private *S*, nucleotide diversity (*π*), Watterson’s estimator (*θ*_W_), Tajima’s *D* (*θ*_D_), relative divergence (*F*_ST_), absolute divergence (*D*_XY_), and site frequencies. In principle any summary statistic can be included, contingent on the data and appropriate additions to the programs’ scripts. To account for potential correlation between summary statistics and to retain only their informative components, we apply a Partial Least Squares transformation (Wegmann et al., 2009).

#### ABC Inference

The estimation of demographic parameters is performed with ABCtoolbox (Wegmann, Leuenberger, Neuenschwander, & Excoffier, 2010), via the ABC-GLM algorithm using the subset of *n* simulations closest to the observed summary statistics. In a first step, LSD infers demographic parameters of putatively neutral loci to obtain the point estimate 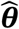. We do so based on simulations of a single locus. In contrast to classic ABC regression approaches, ABC-GLM can readily use such simulations to accurately infer posterior distributions from many loci as it approximates the likelihood function, rather than the posterior distribution (Thalmann et al., 2011). However, we propose a slightly different approach that does not characterize the full posterior distribution, but we found to result in point estimates 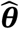 that are more accurate (Supplementary Text S4, Figures S1 and S2) and robust to misidentification of putatively neutral loci (Supplementary Text S5, Figures 4 and S3). Specifically, we first infer locus-specific posterior distributions *π_l_*(***θ***) for each putatively neutral locus with ABC-GLM, then calculate the product of these densities *π*(***θ***) = Π_*l*_ *π_l_*(***θ***), and identify 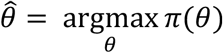.

In a second step, LSD infers locus-specific posterior distributions using ABC-GLM on all loci, either using the same set of simulations as in the first step, or from simulations of a single locus conducted under the parameters 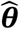, except for the parameters affected by selection (e.g. *M_E_*).

### 2.3 Simulations

To test the performance of the LSD implementation, we simulated pseudo-observed genomes using the program msms under different demographic and selection parameter values, focusing on IM models relevant for the characterisation of local adaptation. We assumed a diploid system, a common mutation rate *μ* = 5×10^−7^ per bp per generation, and all loci to be biallelic with ancestral allele *a* and derived allele *A*. Each simulated pseudogenome comprised *n_n_* = 1,000 neutral loci and *n_s_* = 50 selected loci of 5kb length, for a total (pseudo-genome) size of 5.25Mb.

#### Demography

We simulated four models representing different levels of complexity in terms of population structure and demographic history (Figure 2, Supplementary Text S1). In all models, selection is inferred from the reciprocal scaled migration rates between two contrasting environments (*M*_12_, *M*_21_; where *M_E_* = *Nm_E_*). We used neutral migration rates *M*_12_ = *M*_21_ = *M* = 0.5, 5 and 50 migrants per generation and inferred selection as deviations from these rates. We use model 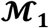 to represent a simplified, generalised model of local adaptation, model 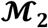 to represent a more complex case of local adaptation comprising multiple, structured populations, model 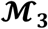 to reflect a scenario typical of glacial-induced secondary contact population dynamics and model 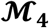 to represent a case of hierarchical divergence with complex demography. The specific parameter choices for all models are given in Supplementary Text S1.

**Figure 2.**
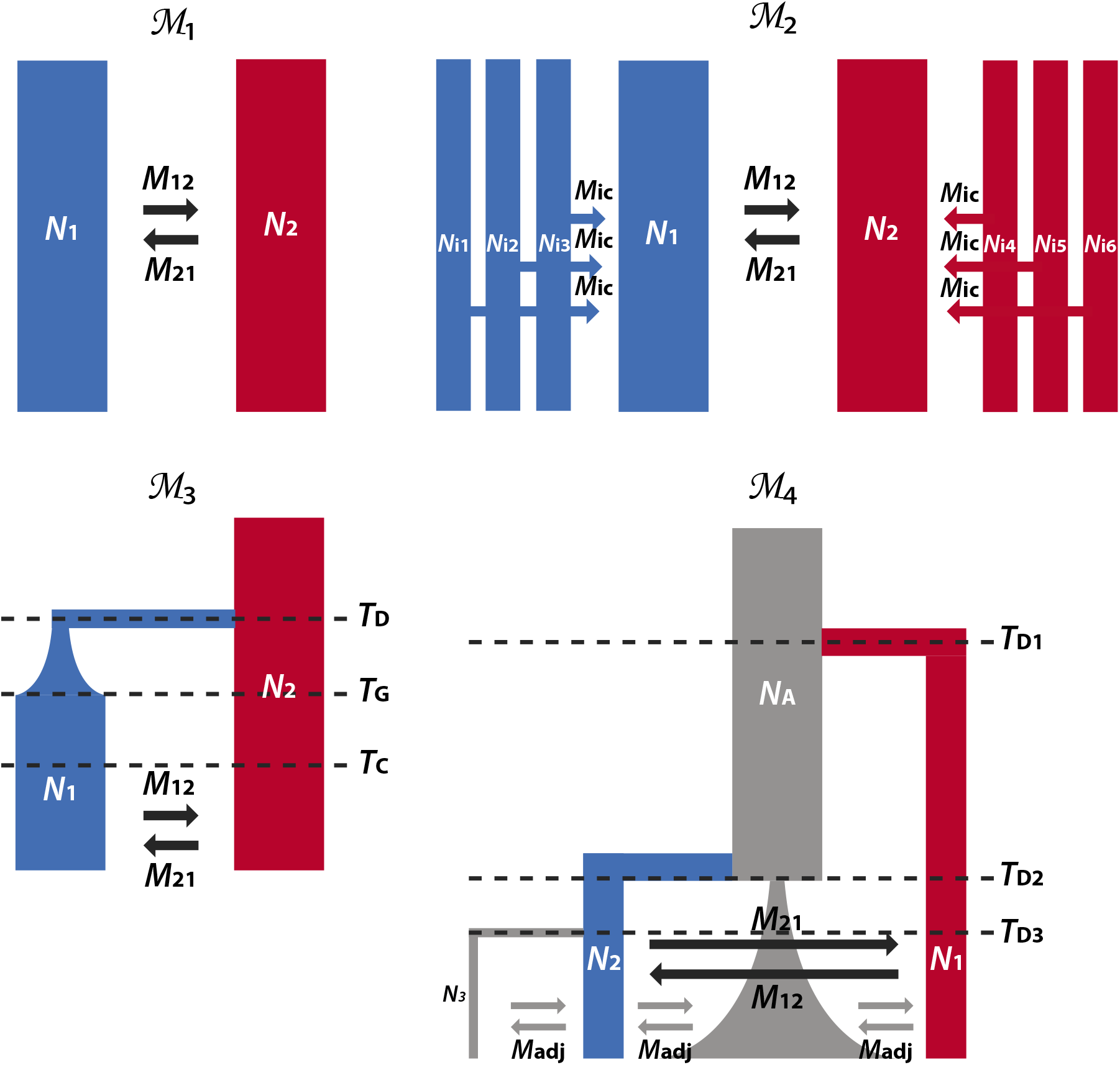
Models used in the simulations and case study. Model 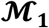 represents a simple 2-deme isolation-with-migration (IM) model with reciprocal migration. Model 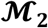 represents a 6-deme island-continent model where common differences between environments are modelled by connecting the sampled demes (i.e. islands) to respective meta-population continents via gene flow. Model 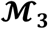 represents a 2-deme divergence with bottleneck and exponential growth model. Model 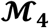 represents a 4-deme hierarchical divergence model with sequential founder events, bottleneck and exponential growth. Red and blue demes reflect contrasting environments, while grey demes reflect neutral environments where no selection acts. In all models, selection is inferred from the deviation from neutrality of the reciprocal migration rates between the two contrasting environments (*M*_12_, *M*_21_).

#### Selection

To simulate genetic trade-offs, selection was simulated on alternate alleles in the contrasting environments, on top of the demographic model. Specifically, we assumed the beneficial alleles to be dominant such that the relative fitness was 1 + *s*_1_ 1,1 and 1, 1, 1 + *s*_2_ for the three genotypes *AA*, *Aa* and *aa* in the demes or meta-populations occupying the two environments, respectively. For the selection coefficients *s*_1_ and *s*_2_, we used all combinations of 0, 0.001, 0.01 and 0.1 and thus included cases of conditional neutrality (CN), in which either *s*_1_ > 0, *s*_2_ = 0 or *s*_1_ = 0, *s*_2_ > 0, as well as cases of antagonistic pleiotropy (AP) with *s*_1_ > 0, *s*_2_ > 0. CN regimes are by definition always asymmetric, while AP regimes can be either symmetric (*s*_1_ = *s*_2_) and asymmetric (*s*_1_ ≠ *s*_2_). We further varied the time of the onset of selection from *T_s_* = 400, 4,000, 40,000 and 400,000 generations ago.

For all models, we considered selection on standing variation with the initial frequency of the derived allele at *f*_1_ = *f*_2_ = 0.1 in all demes. For model 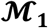, we additionally investigated the case of *de-novo* mutations with initial frequencies *f*_1_ = ½ *N*_1_ and *f*_2_ = 0. These two cases represent the often-considered starting points for local adaptation (Peter, Huerta-Sanchez, & Nielsen, 2012). Depending on the selection regime and due to the stochasticity of drift, the derived allele *A* may sometimes be lost and hence be absent in the simulation of selected loci (especially in the *de-novo* case). Because such a scenario contains no signal for detection of selection, we excluded such simulations (via the-SFC parameter in msms).

#### Assessing accuracy

We inferred selection by contrasting the locus-specific migration rates *M*_12_ and *M*_21_ against their neutral estimates 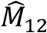 and 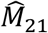 (Figure 3). We evaluated the performance of our LSD implementation at identifying selected loci under these simulations by plotting the true positive rate (TPR) against the false positive rate (FPR) under the choice of HPDI thresholds from 0 to 1, and reporting the area under the curve (AUC) of the resultant receiver operating characteristic (ROC) curve (approximated by the Mann-Whitney U test (Delong & Carolina, 1988)). An AUC value of 0.5 reflects random assignment while that of 1 reflects perfect classification. To evaluate the accuracy of the inferred symmetry of the joint posterior (of reciprocal migration rates), we compared it to the true underlying selection coefficients, under the expectation that deviations from symmetry in the joint posterior should reflect asymmetry in selection regimes. Specifically, we determined for each locus *l* the posterior mass

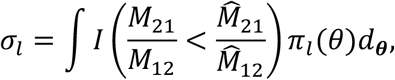

where the indicator function *I*(·) limits the integral to cases in which the deviation of one of the migration rates has reduced more than the reciprocal migration rate compared to a proportional deviation of both migration rates from their neutral estimates 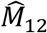 and 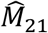. From this, we calculate the asymmetry as

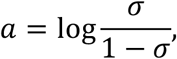

where 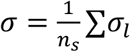 across loci simulated under selection.

AUC and asymmetry are reported for each simulated pseudo-genome, each representative of a unique demographic-selection regime.

**Figure 3.**
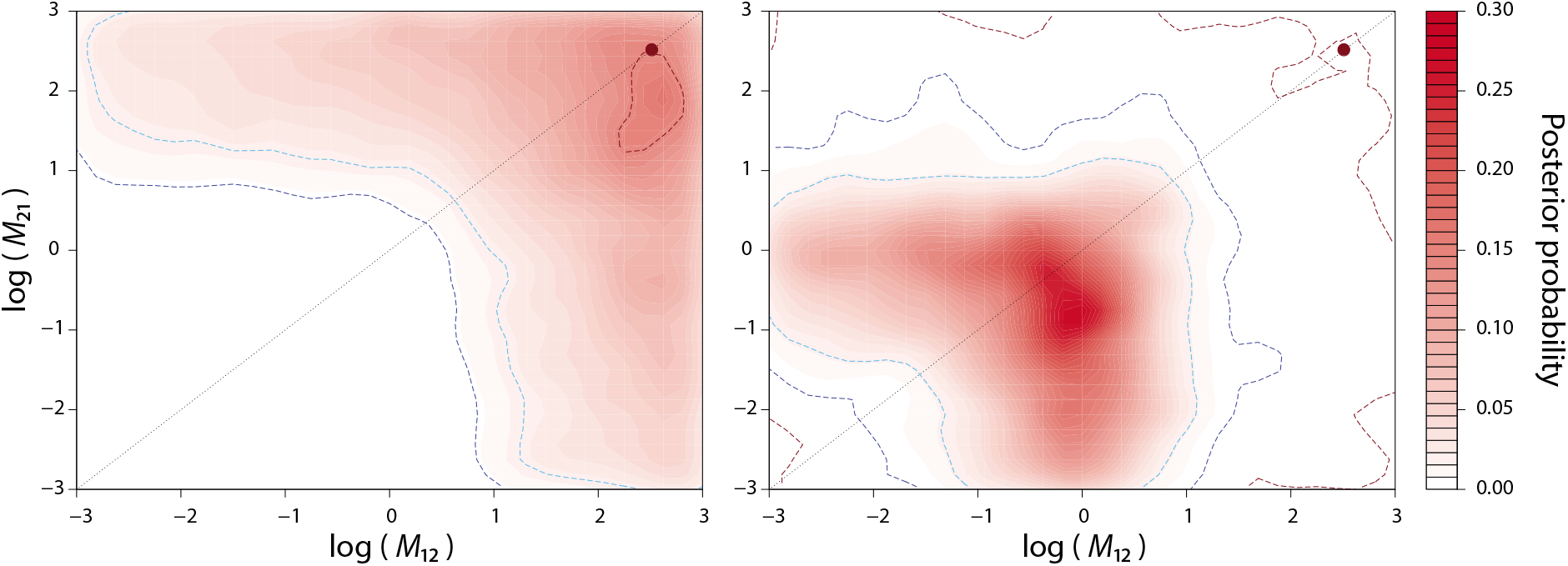
Exemplary joint posterior distribution of reciprocal migration parameters, *M*_12_ and *M*_21_. The neutral joint parameter estimate, as informed by the global posterior distribution of all neutral regions (Figure S1), is indicated by the red dot in the top right corner. The red contours represent the joint posterior distribution of a genomic region (i.e. window), with the blue contours representing the 95% (light blue) and 99% (dark blue) highest density region (HDR) credible intervals. Left – a window not significantly divergent from the neutral estimate; right – a window significantly divergent from the neutral estimate, and with slightly higher relative reduction in *M*_12_ than in *M*_21_.

#### ABC parametrization

Migration rates were drawn from *log*_10_ *M*_12_, *log*_10_ *M*_21_ ~ *U*[−4,3] in all cases, while all other parameters were fixed to their true values (“fixed” parametrisation). We do this to assess the sensitivity and accuracy of LSD under ideal conditions, and to not confound it with model and parameter misspecifications. For ABC parameter estimation, we retain the 1% closest simulations of 250,000 total simulations.

#### Comparison with other methods

We compared the performance of LSD against that of two widely employed genome scan methods, on the simulated pseudo-genomes. pcadapt (Luu et al., 2017) identifies candidate SNPs as outliers with respect to population structure, ascertained via principal component analysis, while OutFLANK (Whitlock & Lotterhos, 2015) infers candidate SNPs by testing against a null model inferred from a highly revised Lewontin-Krakauer model extended to account for non-independent sampling of populations and sampling errors. Given that these approaches are SNP-based (whereas LSD is window-based), we consider a specific genomic window to be an outlier if at least one SNP within that window is called significant. This may reduce false negatives at the expense of inflating false positives, but reflects typical usage of such genome scans. For both methods, the true number of simulated populations was specified and SNPs were thinned to accommodate the particular LD structure of our simulated pseudo-genomes when computing PCs and calibrating the *F*_ST_ null distribution, before running on the full SNP dataset.

For a more realistic comparison where model parameters may not be known with confidence, we also considered cases of 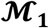 and 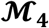 in which all demographic parameters were unknown (“free” parametrisation) and drawn from large, uniform priors *log*_10_ *N_i_* ~ *U*[2,7], *i* = 1,2 for 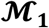 and *log*_10_ *N_i_* ~ *U*[1.5,5], *i* = 1,…,4, *log*_10_ *T_Dj_* ~ [−0.5,1], *j* = 1,2,3, *log*_10_*α* ~ *U*[0,*1*] and *log*_10_ *M_adj_* ~ *U*[0,1] for model 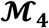. Priors for *M*_12_, *M*_21_ remained the same as before. We used the closest 10,000 out of 1,000,000 simulations to both infer 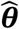 and to identify outlier loci. For added realism, we inferred 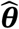 from a full pseudogenome of 1,050 loci, of which 1,000 were neutral but 50 were mis-specified and weakly selected with *s*_1_, *s*_2_ = 0.001; *T_s_* = 4,000, *M_12_* = *M_21_* = *M*.

### 2.4 Case study

To evaluate the performance of LSD on real data, we applied it to the detection of loci underlying floral guides in two parapatric subspecies of *Antirrhinum majus. A. majus* is an herbaceous, perennial, flowering plant native to the western Mediterranean. Owing to its diploid inheritance, relatively short generation time, ability for both self- and crosspollination and rich and varied flower morphology, *A. majus* has lent itself as a model organism for over a century, with several key floral genes being first identified within this genus (Schwarz-Sommer, Davies, & Hudson, 2003; Schwinn et al., 2006). Two subspecies, *A. m. striatum* and *A. m. pseudomajus*, differ in the flower colouration that signposts the pollinator entry point, and form a natural hybrid zone in the Pyrenees that constitutes a benchmark example of divergent selection (Whibley et al., 2006). Several genetic loci have been shown to control the differences in these floral patterns (Bradley et al., 2017; Schwinn et al., 2006), and recently, Tavares et al. (2018) produced evidence of genomic signatures of selection at the ROS and EL loci, of which the former was functionally validated. Here, we apply LSD to sequencing data from this study to isolate the ROS and EL loci and to characterise their underlying selection signal. We filtered the data as in the original study, but mapped on a more recent version of the *A. majus* reference (version 3.0; M. Li et al., 2019).

We modelled this study system via a simple representation (model 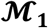) of one population on either side of the hybrid zone (YP1 (*A. m. striatum*) vs MP2 (*A. m. pseudomajus*); populations 2.5km apart) and via a more inclusive island-continent model (model 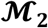) comprising three (distant) populations each per subspecies (CAM, ML, YP1 (*A. m. striatum*) vs MP2, CHI, CIN (*A. m. pseudomajus*); Figure S4), using an estimate of *μ* = 1.7 × 10^−8^ per bp per generation (Tavares et al., 2018) and allowing all *N* and *M* parameters to be free with priors *log*_10_ *N_j_* ~ *U*[1,7], *j* = 1,2, *log*_10_ *M*_12_, *log*_10_ *M*_21_ ~ *U*[−4,4] for 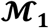 and *log*_10_*N_j_* ~ *U*[3,7], *j* = 1,2, *log*_10_ *N_ik_* ~ *U*[1,6], *k* = 1,…,6, *log*_10_ *M*_12_, *log*_10_ *M*_21_ ~ *U*[−4,4] and *log*_10_*M_ic_* ~ *U*[0,5] for 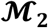. In line with the available pool-seq data for this study, we simulated pooling of individuals in-silico by pooling twice the amount of msms coalescent (haploid) samples as (diploid) individuals in the pooled populations via *LSD-High*. Samples were drawn from a parametric (negative binomial) distribution fitted to the empirical coverage distribution using the ‘*fitdistrplus*’ package (Delignette-Muller & Dutang, 2015) and *LSD-High*. We focused our analysis on chromosome 6 on which the ROS and EL loci lie. To acquire empirical estimates of neutral demographic parameters, we excluded all genomic regions present in the structural annotation plus 10kb flanking regions to generate a subset of putatively neutral regions on that chromosome. 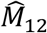 and 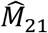 were then estimated via 10kb windows from these neutral regions by retaining the closest 10,000 out of 1,000,000 simulations, as outlined above. To identify selected loci in the second step, we used sliding windows of size 10kb and a 1kb step-size, and retained the closest 5,000 out of the same 1,000,000 simulations.

## 3 RESULTS

### 3.1 Two-deme IM case (model 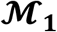)

#### Power to identify selected loci

While our LSD implementation exhibited conservative *p_l_* values (Figure S5), it demonstrated a high diagnostic ability to discriminate between neutral and selected loci (AUC > 0.8) across a large range of migration-selection regimes (Figure 5). Notably, our results point towards an optimal, intermediate rate of migration (*M* = 5) at which selection is best detectable with high AUC values across a large set of selection coefficients. As migration rates increase (*M* = 50), migration from the foreign deme where selection acts on the alternate allele increasingly inhibits the build-up of beneficial polymorphisms in the local deme, in which case the power to detect selected loci becomes limited to scenarios under longer regimes of strong selection. At lower migration rates (*M* = 0.5), long regimes of selection permit the detection of loci under the lowest selection coefficients, but power decreases for younger times compared to scenarios simulated under intermediate migration rates. This owes to LSD relying on the reduction of effective migration relative to neutral or genome-wide expectations, which in this case is already at a low level.

The power to detect selection increased with increasing selection coefficients when these were similar (*s*_1_ ≈ *s*_2_, cells along diagonal of sub-panels in Figure 5). In such cases, stronger selection coefficients on alternate alleles increasingly polarise and ultimately maintain larger allele frequency differences between the two environments. In tandem, the power to detect selection also generally increased with the time since the onset of selection *T_s_*. In contrast, when *s*_1_ ≫ *s*_2_ or *s*_1_ ≪ *s*_2_, one of the two alleles may proceed to fixation, in which case the power to detect selection decays or is lost (e.g. grey cells in left-most column of subpanels when derived allele *A* is lost, and AUC values and inferred (a)symmetries tending toward 0.5 and 0 respectively in bottom row cells when ancestral allele *a* is lost; Figure 5). This is particularly evident when the onset of selection is more distant in the past.

#### Power to characterise (a)symmetry

A benefit of LSD over classic outlier approaches is that it can provide insight into genetic trade-offs underlying local adaptation, by identifying cases in which selection acts at equal strength in the two demes or metapopulations (symmetric AP), or whether selection coefficients differ considerably (CN or asymmetric AP). As shown in Figure 6, the inferred (a)symmetry generally reflected the true (a)symmetry of the underlying selection coefficients well, particularly for regimes with high power to correctly identify selected loci (Figure 5). In lower powered regimes, we observe some cases where the inferred asymmetry does not reflect the underlying asymmetry of the selection coefficients accurately (e.g. blue cells along diagonals in sub-panels at *T_s_* = 4,000; Figures 6 and S6B). We interpret these results further in the discussion.

#### Standing variation vs *de-novo*

A lower initial frequency of the derived allele may be expected to affect LSD’s power to identify selected loci and its power to capture the underlying (a)symmetry of selection coefficients. However, we find that results for simulations building on selection from the *de-novo* and standing variation cases showed generally very similar patterns (Figures 5, 6 and S6). One notable exception however was the inaccurate inference of (a)symmetry in a few regimes with high power (AUC > 0.8) in the *de-novo* case (e.g. blue cells along diagonals in sub-panels at *T_s_* = 4,000 in Figure S6B). This we attribute to the lower initial frequency of the derived allele *A* and consequently longer time needed to approach drift-migration-selection equilibrium for the *de-novo* cases. This is explored further in the discussion.

### 3.2 More complex cases (models 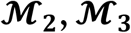 and 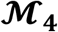)

A key feature of LSD is its potential to explicitly accommodate complex demographies, which can lead to an inflation in false positives when not properly accounted for (De Villemereuil, Frichot, Bazin, François, & Gaggiotti, 2014; Foll & Gaggiotti, 2008; Lotterhos & Whitlock, 2014). Despite the added complexity of models 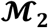 and 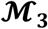, results were generally very similar to that of model 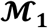, with high power to identify selected loci (AUC > 0.8) across a large range of migration-selection regimes, an optimal migration rate at an intermediate value (*M* = 5), a similar dependence of power to detect selection on *s*_1_, *s*_2_ and *T_s_*, and inferences of (a)symmetry that reflected well the underlying (a)symmetry of selection coefficients (Figures 7, S7 and S8). One notable difference between these models was that model 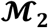 generally required longer time to generate power to detect selection when compared to models 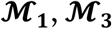, and 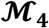, which we attribute to the larger meta-population *N_E_* in model 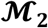. For model 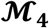, selection coefficients needed to be slightly higher (*s*_1_, *s*_2_ ≥ 0.01; Figure S8) to attain high power to identify selected loci, and to accurately identify asymmetry.

### 3.3 Robustness to mis-specification of the neutral set

As shown for all models and a subset of the regimes (*M* = 5; *T_s_* = 40,000; *s*_1_ = *s*_2_ = 0.01, 0.1), LSD is highly robust to the inclusion of selected loci in the neutral set, with negligible reduction in power (AUC) at up to a 13% mis-specification for model 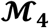 and up to 20% for models 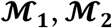 and 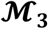 (Supplementary Text S5, Figure 4).

**Figure 4.**
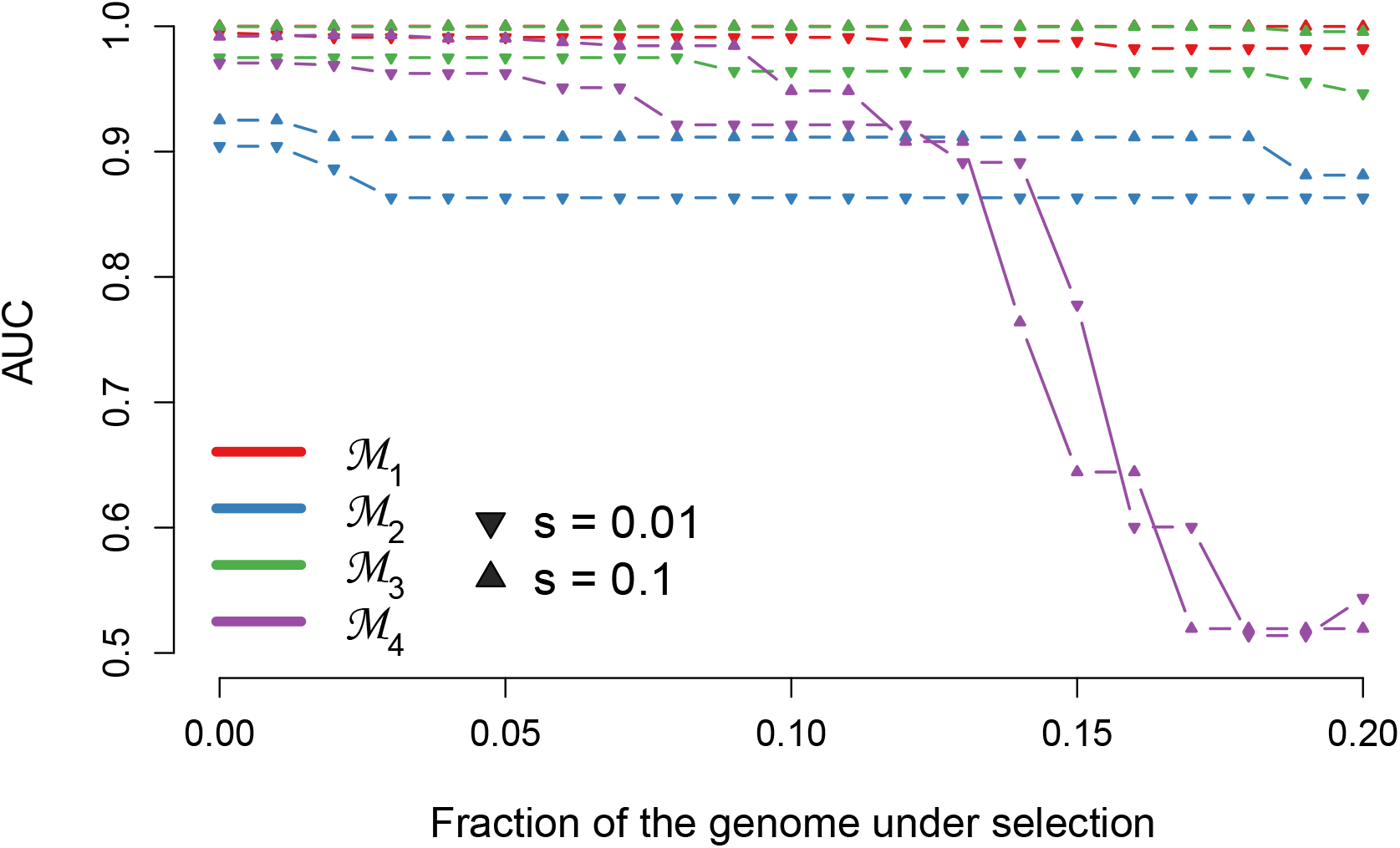
Effect of increasing fraction of mis-specified (aka selected) windows among the neutral set on LSD’s power to detect selection, for all 4 models and intermediate and high selection coefficients (*s*_1_ = *s*_2_ = *s* = 0.01, 0.1; *T_s_* = 40,000), and under neutral migration rates *M*_12_, *M*_21_ = 5. Here the neutral set of 1000 loci comprise a fraction *f* of selected loci and a fraction 1-*f* neutral loci, with 0.0 ≤ *f* ≤ 0.2. The selected loci comprising the pseudo-genome under scan are under the same selection regime as those included as mis-specifications in the neutral set.

### 3.4 Comparison to other methods

The performance of LSD is comparable to that of pcadapt and OutFLANK under model 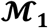, with minimal difference between the methods across the range of migration-selection regimes (Figures 8 and S9). For the complex model 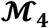 however, both pcadapt and OutFLANK have little to no power to correctly identify selected loci (AUCs ~ 0.5; Figures 8 and S10), while LSD exhibits high AUC scores (> 0.8) in similar migration-selection regimes as it does under models 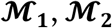 and 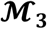. Importantly, the power of LSD to identify loci under selection was very similar (Figures 5, 8, S8A, S9A and S10A) when fixing 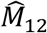 and 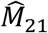 and all demographic parameters not affected by selection to their true value (“fixed” parametrisation), as when estimating 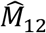 and 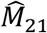 (Figures S11 and S12) and keeping all other parameters free (“free” parametrisation), implying that LSD is robust to parameter specification.

**Figure 5.**
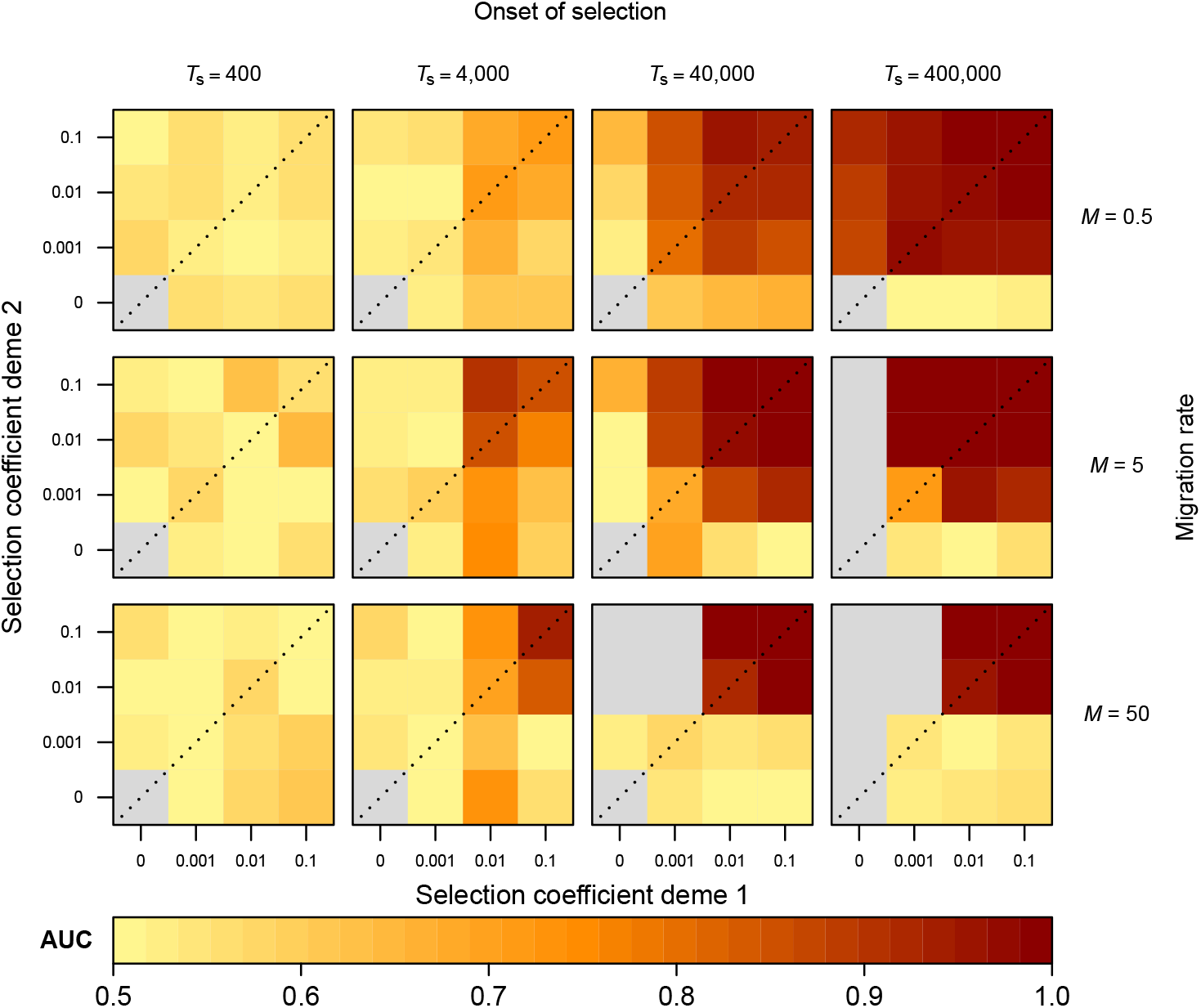
Simulation results showing the effect of migration rate, time of onset of selection and deme-specific selection coefficients on LSD diagnostic performance (AUC), for the 2-deme IM model (model 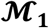; standing genetic variation case). Each cell represents a pseudo-genome simulated under a specific selection regime. The cell colours reflect the AUC calculated by the correct discrimination of 1000 neutral loci and 50 selected loci in the 1050 loci simulated pseudo-genomes. Grey cells indicate selection regimes where the derived allele is always lost.

### 3.5 Case study results

We identified a region of reduced effective migration between 52.9-53.2MB on chromosome 6 (Figure 9), consistent with the location of the ROS and EL loci (Tavares et al., 2018). Under model 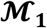, this region is characterised by a set of smaller, multiple peaks (*p_l_* < 0.01) reflecting signatures identified by previous authors, with the left-most peaks corresponding to ROS1 and ROS2 (shaded in red) and the right peaks to EL (shaded in green; Figure 9B). The joint posterior probability distributions reveal symmetric selection acting on both regions, implying that selection acts with similar strength in the two populations. Under Model 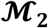, we find fewer outliers in the ROS-EL region than in model 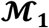, with the left-most peak in this region corresponding to ROS2 and the right peaks consistent with EL. ROS1 appears to be less of an outlier than in model 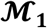(*p_l_* ≈ 0.02). In contrast to model 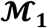, the ROS2 and EL peaks in model 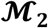 are characterised by asymmetry, specifically with stronger selection acting in the populations of *A. m. pseudomajus* than in the populations of *A. m. striatum*.

## 4 DISCUSSION

The trajectory of selected loci depends on demographic and selection parameters that define the system, namely the effective population sizes, effective migration rates and selection coefficients, as well as the intrinsic properties of mutation and recombination. Despite well-developed theory which relates the effect of population parameters on the trajectory of selected alleles, few methods or empirical studies have combined estimates of differential selection with explicit quantification of migration rates and effective population sizes to examine the conditions under which local adaptation can arise. Here, we introduce such a method, LSD, which identifies candidate loci based on divergent population parameters using explicit demographic models, and demonstrate that under certain demographic-selection regimes, it can both detect and elucidate the processes underlying signatures of selection. While LSD is flexible regarding the choice of demographic models employed and can apply to single and multiple populations, we focus here specifically on processes that lead to selection against gene flow, namely local adaptation and extrinsic reproductive barriers, that can be inferred via their expectation to reduce *M_E_*.

### 4.1 Identifying selection

Using simulations, we demonstrate that LSD has high diagnostic power (AUC > 0.8) to identify selected loci across a large range of demographic-selection regimes. This power relies upon two fundamental aspects that contribute to generating observable patterns. First, selection must effectively be realised, i.e. result in a frequency shift of the beneficial allele. This requires that the strength of selection and initial frequency of the beneficial allele be sufficient to both counter the homogenising effect of migration (Felsenstein, 1976; Haldane, 1930; Lenormand, 2002; Olson-Manning, Wagner, & Mitchell-Olds, 2012; M. Slatkin, 1973; Yeaman, 2015) and the eroding effect of drift (Wright, 1931). Secondly, the genomic data must contain signatures of selection that can be detected. In the case of LSD, this requires that the signatures of selection are discernible from the underlying noise (drift and migration) that characterises the system, which demands sufficient time for said signatures to be reflected in the employed statistics and hence in the inferred parameters *N_E_* or *M_E_*. A lack of power in LSD must be interpreted considering these two conceptually different perspectives. Notably, the lack of discrimination power for high migration rates and low selection coefficients can be attributed to selection failing to realise as a consequence of local, beneficial alleles being swamped by immigrant, maladaptive alleles. In contrast, the lack of signal under low migration rates constitutes a methodological limitation of our implemented model, as it becomes increasingly difficult to detect reductions in effective migration when neutral or genome-wide migration rates are already at a low level; even when selection is effectively being realised in the demes. This is analogous in effect to the loss of power to detect selection in highly differentiated populations in *F*_ST_ outlier tests (Hoban et al., 2016; Martin et al., 2013). Under the same principle, we argue that the converse expectation can be assumed to hold for loci underlying adaptive introgression or balancing selection. That is, we expect power to detect such loci to be low when populations are minimally differentiated and high in highly divergent systems, as the detection of candidate loci in these cases is informed by increased effective migration.

The power of LSD to correctly identify selected loci generally increases with stronger selection coefficients and longer time since the onset of selection, though with exceptions. Specifically, if selection is of similar or equal strength in both demes or meta-populations, we observe a strong correlation between the power to detect selection and the true underlying selection coefficients. This follows theory which states that the reduction in effective migration is proportional to the strength of selection (Petry, 1983). However, we defer from translating these changes to explicit selection coefficients because in addition to the strength of selection, changes in effective migration are also a function of the recombination rate between linked and selected loci (Cutter & Payseur, 2013; Lotterhos, 2019; Petry, 1983). If selection differs strongly between demes or meta-populations (*s_i_* ≫ *s_j_*), or when the onset of selection is sufficiently distant in the past, however, one allele may become fixed in the system. In such a case, the signal to detect selection rapidly decays (Huber, DeGiorgio, Hellmann, & Nielsen, 2016; Przeworski, 2002). Moreover, we observed little power to detect very recent selection, intrinsically related to our choice of summary statistics (Hohenlohe, Phillips, & Cresko, 2010). That said, extending LSD to include additional statistics sensitive to linkage disequilibrium such as extended haplotype homozygosity (EHH) (Sabeti et al., 2002) or single density score (SDS) (Field et al., 2016) will increase the power to detect more recent selection. Finally, given that LSD relies on localised deviations in effective demographic parameters, we predict that it may miss signals of polygenic selection, as signals of selection become increasingly hard to distinguish from the genomic background the smaller the locus effect sizes are.

From our simulations, we find that the power to detect selection is similar between the *de-novo* and standing genetic variation regimes (model 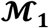; Figures 5 and S6A), likely as a result of only considering loci for which the derived allele was not lost. This particularly affected the *de-novo* case, under which the derived allele was lost in most simulations. Indeed, this is in line with theoretical expectation (Olson-Manning et al., 2012) and supports the notion that most empirical cases of local adaptation attribute selection of advantageous alleles to arise from standing variation (Jones et al., 2012; Lai et al., 2019; Reid et al., 2016). To clearly distinguish between these regimes, additional information on the evolutionary history of the system such as allele age, mutation rate or supplementary phylogenetic information is required (Peter et al., 2012).

### 4.2 Comparison to existing methods

LSD’s power to detect selection is comparable with that of pcadapt and OutFLANK under the simple model 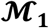, but strongly outperform these alternatives under the more complex model 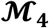. These results appear to hold regardless of whether prior knowledge of model parameters is confidently known for LSD, as LSD exhibits similar performance under the ideal “fixed” and realistic “free” parametrisations. This suggests that the methods pcadapt and OutFLANK utilise to address population structure, namely PCA and an implicitly modelled *F*_ST_ null distribution, respectively, are sufficient to control for the effects of relatively simple demography, but insufficient to capture more complex demographies. By modelling complex demographies explicitly, LSD is less affected by model complexity, though this extra power comes with some costs.

Firstly, a demographic model needs to be specified that appropriately describes the neutral genetic variation of the system, allows for inferences of selection through changes in demographic parameters (e.g. *N_E_* or *M_E_*), and is sufficiently simple to remain computationally tractable. A preliminary analysis of model choice may therefore constitute a prerequisite to successfully recapitulate the signal of complex evolutionary histories in the simulated data. Importantly, the model should always be validated by demonstrating that the observed data can be accurately and sufficiently captured (Figure S13).

Secondly, LSD remains computationally more demanding than both pcadapt and OutFLANK. Inferring demographic parameters is generally computationally challenging as the underlying genealogies need to be integrated out numerically (Hey & Nielsen, 2007), which for complex models usually requires simulation-based approaches such as ABC. Existing ABC approaches to infer locus-specific parameters (Bazin et al., 2010; Kousathanas et al., 2016) are difficult to scale-up to genome-wide data as they require the simulation of very many loci. To circumvent this problem, LSD implements an efficient ABC approach that requires simulations of single loci only, which is possible because LSD neither attempts to infer the hierarchical distribution of locus-specific parameters nor to obtain posterior estimates on whether a locus is affected by selection. Instead, LSD identifies loci under selection by quantifying whether locus-specific estimates of demographic parameters are incompatible with those estimated from a set of putatively neutral loci.

The *a priori* identification of this neutral set constitutes the third requirement. Such a set may be informed by the particular structural or functional class the sites belong to (Williamson et al., 2005) and may for instance consist of genomic regions not linked to structural annotations. Alternatively, a more naïve strategy may rely on the whole genome or a random subset of the genome to reflect neutral diversity. Even if this assumption of neutrality is violated (Begun et al., 2007; Fay, Wyckoff, & Wu, 2002; H. Li & Stephan, 2006), we show that our method to estimate 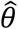 is robust to the mis-identification of neutral loci, with minimal effect on power even at high percentages of up to 20% mis-specification. We attribute this to our method of estimating 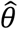 as the product of per-locus posterior densities, which amplifies the signal (density) according to majority rule.

We finally note that most widely applied genome scans methods (e.g. pcadapt and OutFLANK) detect outliers at the SNP level, while our current implementation of LSD focuses on genomic windows. Focusing on genomic windows offers a means to aggregate information across linked loci, thereby potentially increasing power and reducing false positives from spurious signals at individual SNPs (e.g. Galimberti et al., 2020). These benefits are conditional on a choice of window size compatible with the LD decay in the system, as this maximises the power and accuracy to capture the signal generated by selection. The LSD framework is by no means limited to genomic windows however, and can be extended to SNP data when using an appropriate simulator and summary statistics calculator.

### 4.3 Revealing trade-offs underlying selection

LSD can also shed light on the directionality of selection by inferring the (a)symmetry in deviations of migration rates between populations. In our simulations, this inferred (a)symmetry accurately reflects the (a)symmetry in the underlying selection coefficients for older onsets of selection, but less so for more recent onsets (Figures 6, 7B, S6B, S7B). This is because inferred asymmetries in deviations of effective migration rates are also affected by asymmetries in allele frequencies of the beneficial allele. For instance, a new beneficial mutation that arises in one of two demes will initially be rare among migrants. Only as its frequency increases will selection start to act against immigrants in both demes (Figure 10). Hence, the inference of asymmetry in LSD may include cases where selection is effectively asymmetric, but also those in which selection is effectively symmetric but prior to driftmigration-selection equilibrium. A direct link between the inferred (a)symmetry in deviations of migration rates and the (a)symmetry in underlying selection coefficients is only established through time. We note that in practice, however, the interpretation of the results is straightforward, as the inference of directionality is only meaningful for loci identified as under selection i.e. with low *p_l_* values. Such loci can only be inferred in regimes with high AUC, which generally preclude regimes characterised by recent onsets of selection. For these (high AUC) regimes, we generally find the estimated (a)symmetries to reflect the true (a)symmetry in selection coefficients accurately. Stated succinctly, LSD’s inferred directionality is generally accurate and meaningful for identified candidates, and potentially inaccurate but irrelevant for other loci.

**Figure 6.**
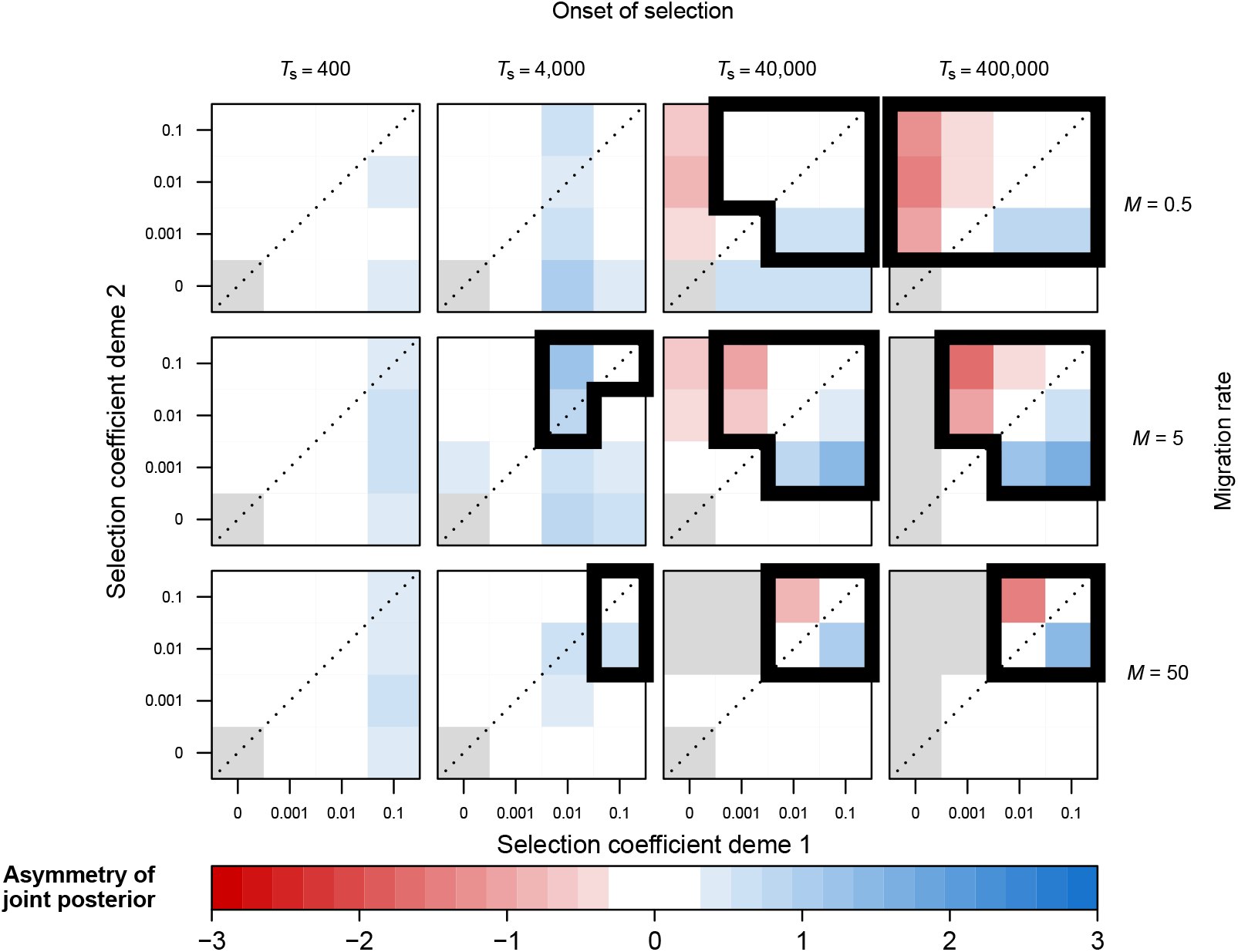
Simulation results showing the effect of migration rate, time of onset of selection and deme-specific selection coefficients on LSD inferred (a)symmetry of selection, for the 2-deme IM model (model 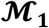; standing genetic variation case). Each cell represents a pseudo-genome simulated under a specific selection regime. The cell colours reflect the (a)symmetry values inferred by LSD, where a value of 0 reflects perfect symmetry of the joint posterior while values divergent from this reflect asymmetry. Cells surrounded by thick lines indicate the values of (a)symmetry for regimes expected to generate meaningful signal (AUC>0.8 in Figure 5). Grey cells indicate selection regimes where the derived allele is always lost.

**Figure 7.**
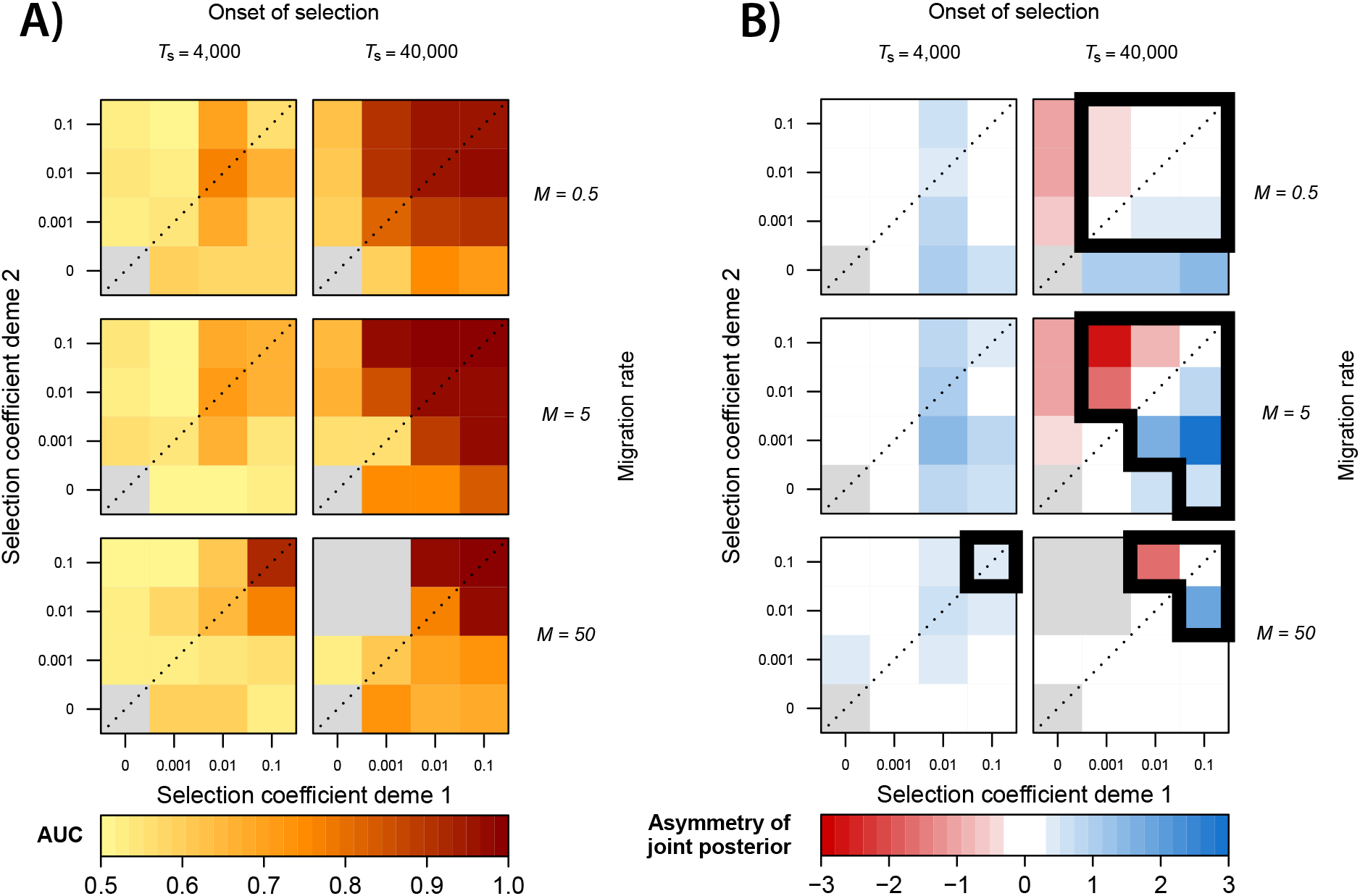
Simulation results for a 2-deme divergence with bottleneck and exponential growth model (model 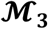; standing genetic variation case) showing the effect of migration rate, time of onset of selection and deme-specific selection coefficients on A) LSD diagnostic performance (AUC) and B) LSD inferred (a)symmetry of selection. Divergence time of the two populations, *T*_D_, is 200,00 generations ago. Each coloured cell represents a pseudogenome simulated under a specific selection regime. Grey cells indicate selection regimes where the derived allele is always lost. B) Cells surrounded by thick lines indicate the values of (a)symmetry for regimes expected to generate meaningful signal (AUC>0.8 in Figure 7A).

**Figure 8.**
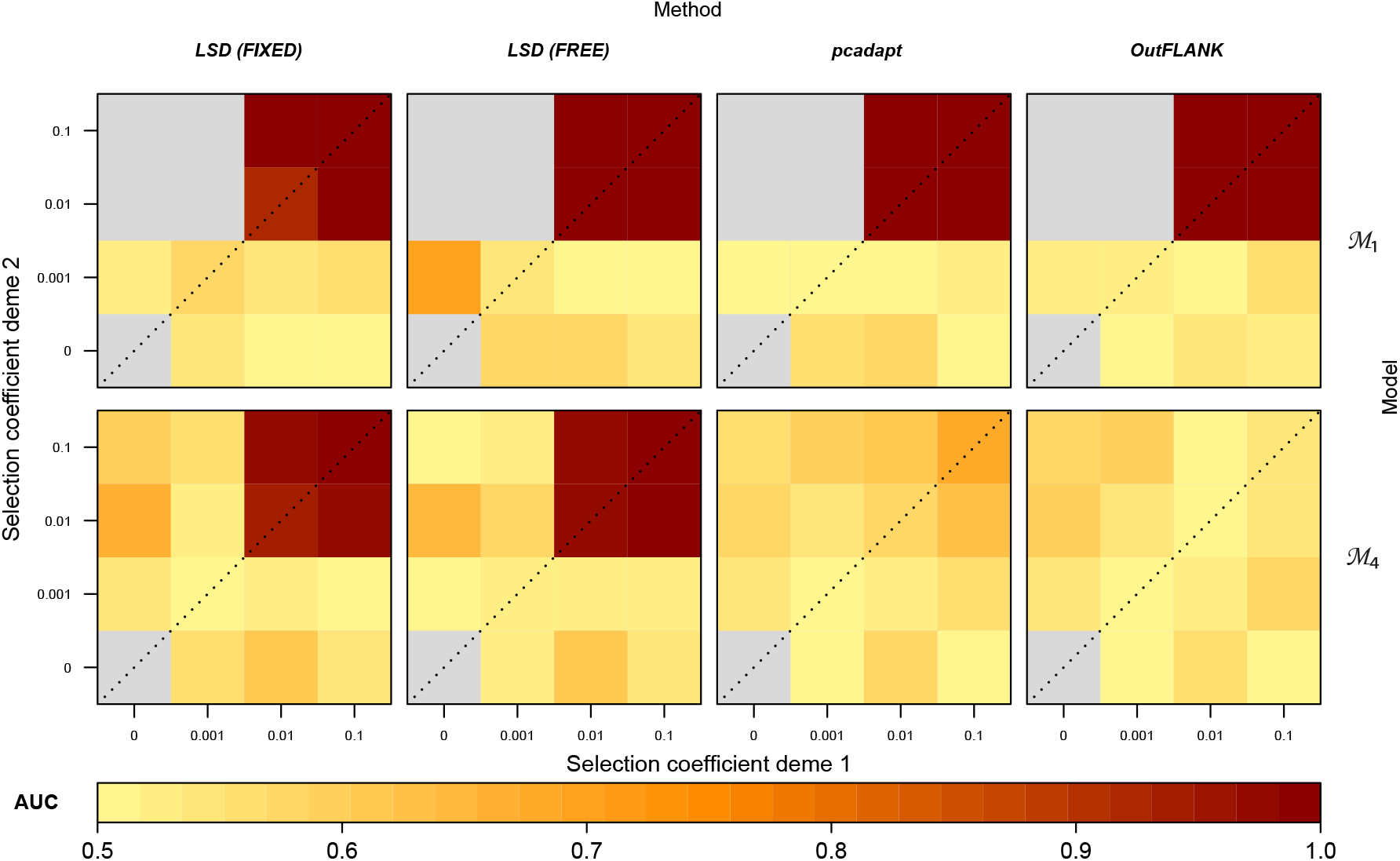
Comparison of power to detect selection (AUC) between LSD, pcadapt and OutFLANK. Simulation results are shown for a simple 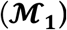 and a complex demographic model 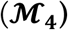 and for a subset of the tested migration-selection regimes at *T_s_* = 40,000 and *M* = 50 (full results in Supplements). For LSD, two parametrisations are performed: i) assuming non-*M* demographic parameters and neutral 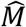 to be fixed to the true values (LSD FIXED) and ii) allowing non-*M* demographic parameters to be drawn from large prior ranges and using neutral 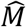 estimated from the pseudo-genomes (LSD FREE). Grey cells indicate selection regimes where the derived allele is always lost.

**Figure 9.**
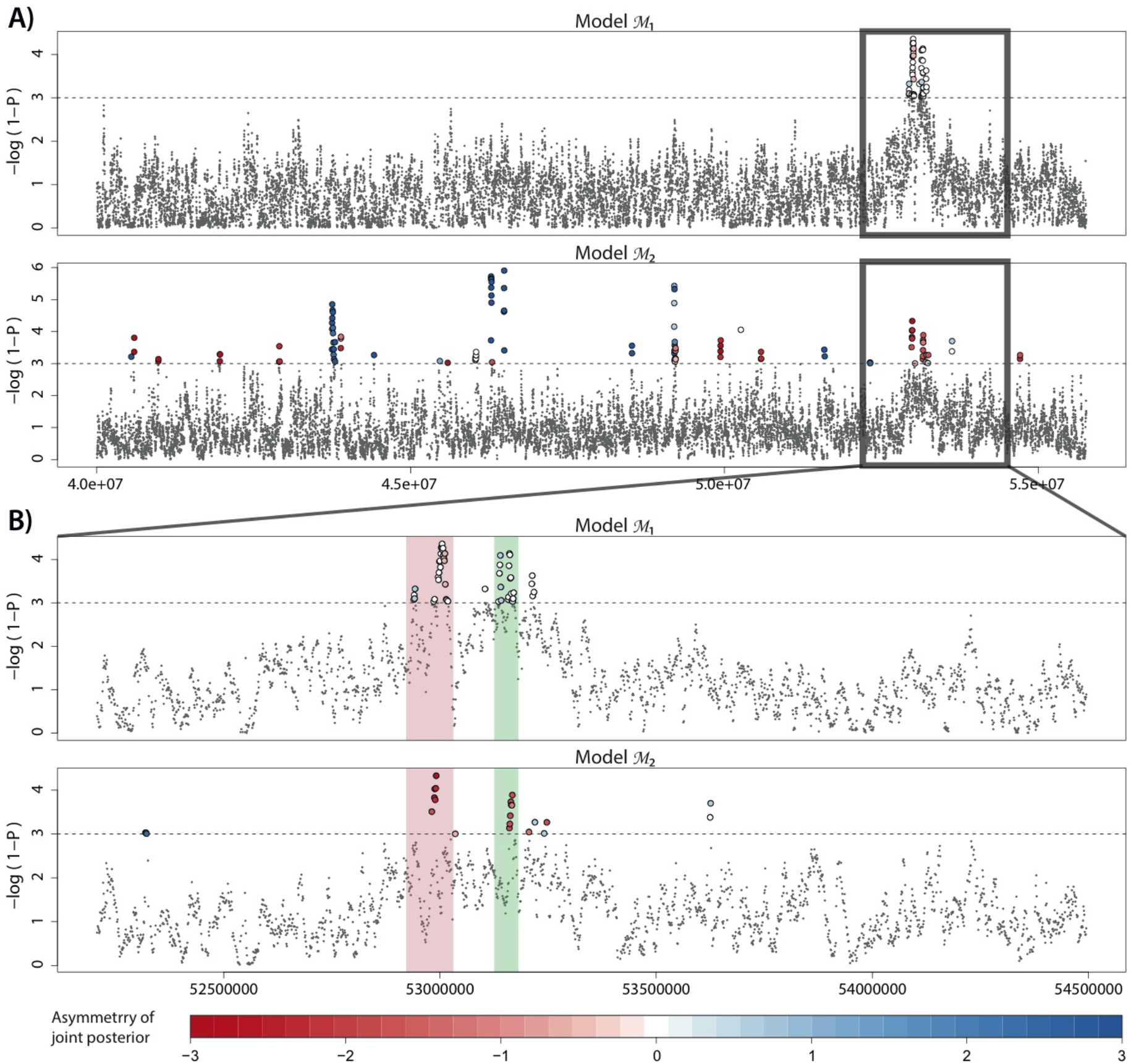
Manhattan plot for the LSD scan of the *A. m. striatum-A. m. pseudomajus* system. The posterior probability of observing the neutral estimate for 10kb windows (1kb step-size) is plotted for A) a 16Mb region of chromosome 6, under models 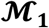 and 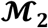, and B) a 2Mb zoomed-in region of chromosome 6 focusing on the ROS-EL region, under the same two models. The horizontal dashed line indicates a 99.9% posterior probability of deviating from neutral expectations. Colour for loci above this threshold denotes the joint (*M*_12_, *M*_21_) posterior (a)symmetry, and reflects the relative strengths of selection in the two divergent demes or subspecies. A large divergent peak centred around the ROS-EL region (A) is composed of a set of smaller peaks (B), consistent with the ROS (red) and EL (green) loci.

**Figure 10.**
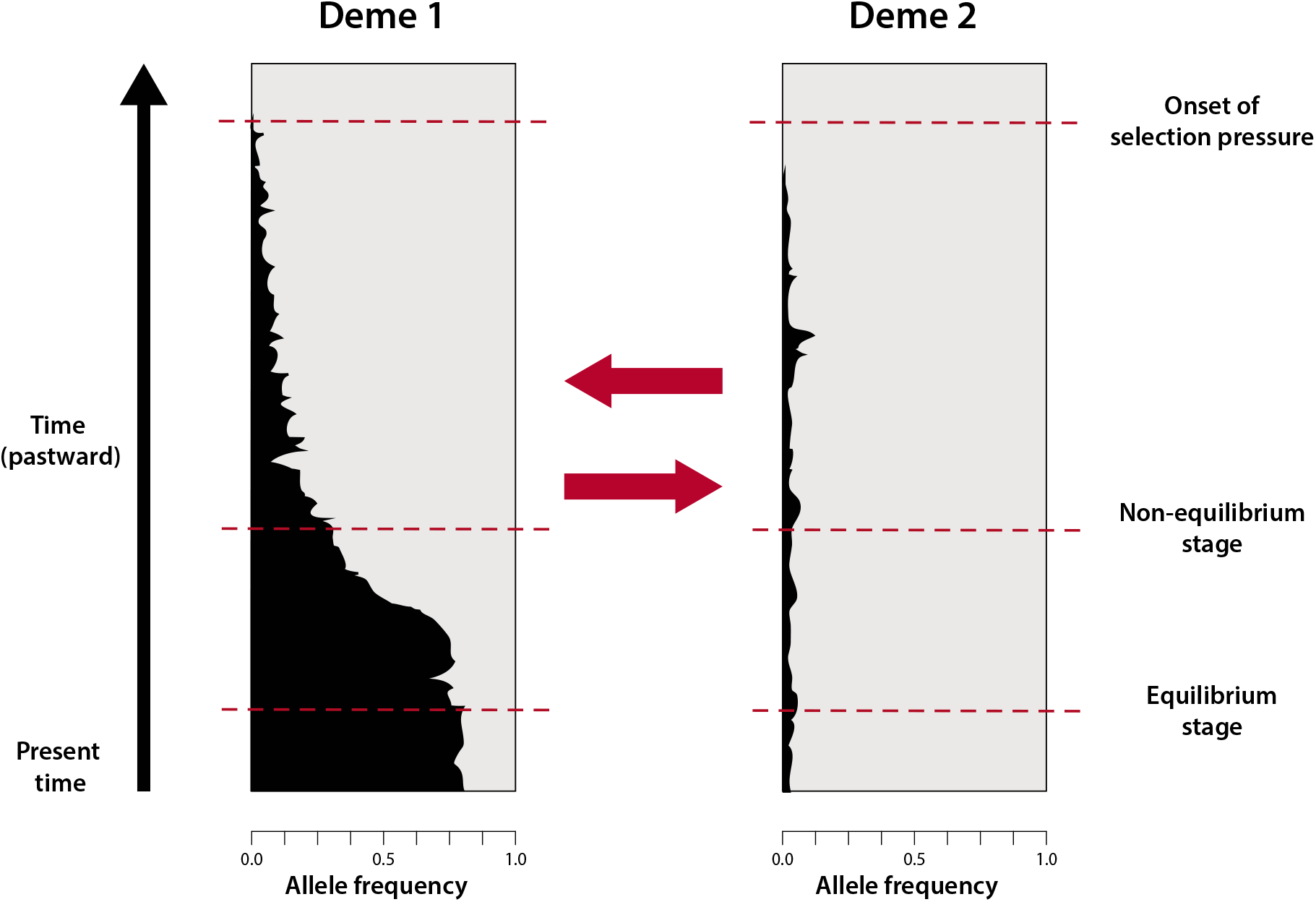
Conceptual illustration of allele frequency trajectories over time in a 2-deme IM model 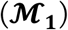, for an example *de-novo* case and antagonistic pleiotropic selection regime. The frequency of derived allele *A* is indicated in black and that of ancestral allele *a* in grey. Red arrows represent migration. Prior to reaching drift-migration-selection equilibrium, estimated asymmetries in effective migration rates are also affected by asymmetry in allele frequencies.

The ability of LSD to infer the directionality of selection directly from genomic data can greatly facilitate investigations of genetic trade-offs underlying adaptation, which are seldom performed due to the considerable effort required to set up field trials of recombinant lines. As shown above, the inference of symmetry in LSD-identified candidates accurately reflects cases of AP with equal strength of selection on alternate alleles in the contrasting environments. The inference of asymmetry on the other hand can either indicate AP with stronger selection in one environment than the other, or CN. From our simulations, we find that scenarios reflecting AP are generally more readily detected than those reflecting CN. Given that selection acts only upon one of the two alleles in the latter case, fixation becomes likely and the ability to detect selection is transient. This implies that there may be an observation bias between AP and CN; such that the inference of CN may be comparatively under-represented. This bias appears to contrast with that reported in ecological literature, where instances of AP are more rarely detected compared to CN due to the additional power required to statistically prove differential fitness concurrently in two environments (Anderson et al., 2013). LSD may further complement field trials as such experiments typically test genetic trade-offs under contemporary selective environments, which may not reflect past conditions driving the observed adaptive responses, but whose signature may still be inferred from genomic data. Using LSD to formulate expectations about fitness effects and to inform the choice of environmental conditions under which to validate identified candidate genes can thus greatly aid such experiments.

### 4.3 Real-world application

We demonstrate a real-world application of LSD by successfully isolating and characterising the selection signal of loci underlying an extrinsic reproductive barrier in *A. majus*. Our results from contrasting a single population (model 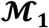) and three populations (model 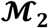) per subspecies both identified the ROS and EL loci which were previously reported to underlie differences in floral patterns between these subspecies (Tavares et al., 2018). Interestingly however, our results characterise selection at these loci as symmetric under model 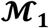 and asymmetric with stronger selection acting on *A. m. pseudomajus* than in *A. m. striatum* under model 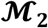. This exemplifies that results of LSD genome scans are conditional on the model and populations used, such that here, model 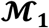 uncovers population-pair specific differences at the contact zone (YP1 vs MP2) while model 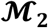 reveals common (global) differences between the two subspecies. We do not necessarily expect these two signals to be identical, and indeed, Tavares et al. (2018) showed different *θ*_W_ and *F*_ST_ estimates between distant and close *A. m. striatum-A. m. pseudomajus* population pairs. Given that there is no evident difference in environment or pollinators on opposite sides of the hybrid zone, reproductive barriers in this system have often been proposed to be maintained through selection against hybrids and frequency-dependent sexual selection mediated by pollinator preference for the dominant flower phenotype on either side of the contact zone. However, whether selection on alternate alleles follows the same positive frequency-dependence across the broader scale including more distant populations is currently unknown. The difference in signal between local pairs at the contact zone 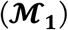 and the global set 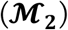 may be generated by different frequency-dependent selection curves for the alternate alleles and potentially loss of AP away from the contact zone (Figure S14).

## 5 CONCLUSION

Loci under selection are predicted to exhibit genealogies with demographic parameters divergent from those of neutral non-linked regions, leading to heterogeneity in demography across the genome. In this study, we condition the identification of candidate loci on divergent population parameters using explicit demographic models, and demonstrate that under certain conditions of migration, selection strength and onset time, we can both detect and elucidate the underlying processes driving signatures of selection. Incorporating and utilising the inference of demographic parameters in the identification of candidate loci address some key issues and assumptions that prevail in the discrimination of selected variants, namely 1) the explicit consideration of demography, 2) heterogeneity in drift and gene flow across the genome, 3) information synthesis of multiple, complementary summary statistics, and 4) transparency towards underlying driving mechanisms.

Our power analysis using simulations shows that LSD, and our implementation of it, represents a powerful method for detecting selection that is robust to different and complex demographies. Furthermore, given that certain demographic parameters, e.g. migration, are not inherently commutative, we show that the directionality or population-specificity in selection can be inferred. This can facilitate identifying in which environment selection acts and hence elucidate genetic trade-offs; bridging an analytical divide between experimental ecology and population genomics. Importantly, the proposed approach as well as our implementation is not limited to the demographic models investigated here, nor the explicit choice of simulation programs or summary statistics used. This flexibility and customisability of LSD can facilitate e.g. more realistic accommodation of recombination (via different coalescent simulators), improved detection of more recent selection (via linkage-informative statistics), and inference of other modes of selection (e.g. balancing selection) and adaptive introgression by conditioning the detection of selection on e.g. increase (rather than reduction) of *M_E_* or changes in *N_E_* relative to neutral expectations.

## ACKNOWLEDGEMENTS

We would like to thank the Genetic Diversity Centre at ETH Zurich and in particular Niklaus Zemp for providing IT support. This work was supported by the Swiss National Science Foundation (SNSF) grants 31003A_160123 and 31003A_182675 awarded to AW and grant 31003A_173062 awarded to DW.

## DATA ACCESSIBILITY

We provide scripts to perform LSD genome scans at the GitHub repository: https://github.com/hirzi/LSD.

## AUTHOR CONTRIBUTIONS

HL, DW, AW and SF designed the study. HL wrote the LSD scripts and performed the simulations and analyses. HL and DW developed the methods. HL wrote the manuscript, which all authors critically revised.

