## Supplementary Material for "Identifying loci under selection via explicit demographic models"

### SUPPLEMENTARY TEXT

#### S1. Model definitions ( $\mathcal{M}_1, \mathcal{M}_2, \mathcal{M}_3$ and $\mathcal{M}_4$)

We simulated four models representing different levels of complexity in terms of population structure and demographic history. In all models, we used neutral migration rates $M = 0.5$, 5 and 50 migrants per generation and inferred selection as deviations from these rates.
For model diagrams, refer to Figure 2.

##### Model $\mathcal{M}_1$

A simple 2-deme IM model with effective population sizes $N_1 = N_2 = 10{,}000$ and symmetric migration rates $M_{12} = M_{21} = M$.

##### Model $\mathcal{M}_2$

A 6-deme IM model comprising 2 contrasting environments of 3 demes each structured as 'islands' with effective population size $N_i = 1{,}000$ connected via migration $M_{ic} = 100$ to meta-population ‘continents’ of size $N_1 = N_2 = 100{,}000$. These continents exchange migrants at symmetric rates $M_{12} = M_{21} = M$.

##### Model $\mathcal{M}_3$

A 2-deme divergence with bottleneck and exponential growth model. Under this model, two demes split from an ancestral population of size $N_A = 10{,}000$ at $T_D = 200{,}000$ generations ago. Following divergence, deme 2 stays constant at $N_2 = 10{,}000$, while deme 1 undergoes a sudden bottleneck, immediately followed by exponential growth with rate $\alpha = 2$ until $T_G = 160{,}000$ generations ago, at which $N_1 = 10{,}000$ is reached and thereafter remains constant. Demes 1 and 2 are initially separated with no gene flow between demes (isolation period), after which secondary contact is established at $T_c = 20{,}000$ generations ago with symmetric migration rates $M_{12} = M_{21} = M$.

##### Model $\mathcal{M}_4$

A 4-deme multiple divergence model with successive founder events and changing population size. Under this model, 2 demes split from a large ancestral population of size $N_A = 10{,}000$ at times $T_{D1} = 160{,}000$ and $T_{D2} = 60{,}000$ generations ago, giving rise to demes $N_1$ and $N_2$ respectively, both of population size 4,000 which thereafter remain constant through time. A subsequent divergence event occurs in deme $N_2$ at time $T_{D3} = 40{,}000$ generations ago, giving rise to deme $N_3$ of population size 400 which thereafter remains constant through time. Each split in this model can be seen as a founder event retaining 10% of the size of the source population. Concurrent with the second divergence event, the ancestral deme experiences a bottleneck followed by exponential population growth of rate $\alpha = 2.5$ reaching a size of 200,000 in the present. Migration is allowed between adjacent demes ($M_{adj} = 2$) and between demes 1 and 2, which are initially isolated post-divergence ($M_{12} = M_{21} = 0$), but establish secondary contact at time $T_{D3} = 40{,}000$ generation ago at symmetric rates $M_{12} = M_{21} = M$.

#### S2. *msms* command lines for generating models $\mathcal{M}_1$, $\mathcal{M}_2$, $\mathcal{M}_3$ and $\mathcal{M}_4$

For all models, we used neutral migration rates $M_{12} = M_{21} = M = 0.5$, 5 and 50 migrants per generation.

**$\mathcal{M}_1$ (neutral demography)**

msms 80 1 -t 10 -I 2 40 40 -n 1 1 -n 2 1 -m 1 2 $M$ -m 2 1 $M$ -N 10000

**$\mathcal{M}_1$ (selection addendum)**

-SI $T_S$ 2 $f_1$ $f_2$ -Sc 0 1 $s_1$ 0 0 -Sc 0 2 0 0 $s_2$ -Sp 0.5 -SFC

**$\mathcal{M}_2$ (neutral demography)**

msms 240 1 -t 10 -I 8 40 40 40 0 0 40 40 40 -n 1 0.1 -n 2 0.1 -n 3 0.1 -n 4 10 -n 5 10 -n 6 0.1 -n 7 0.1 -n 8 0.1 -m 1 4 100 -m 2 4 100 -m 3 4 100 -m 8 5 100 -m 7 5 100 -m 6 5 100 -m 4 5 $M$ -m 5 4 $M$ -N 10000

**$\mathcal{M}_2$ (selection addendum)**

-SI $T_S$ 8 $f_1$ $f_1$ $f_1$ $f_1$ $f_2$ $f_2$ $f_2$ $f_2$ -Sc 0 1 $s_1$ 0 0 -Sc 0 2 $s_1$ 0 0 -Sc 0 3 $s_1$ 0 0 -Sc 0 4 $s_1$ 0 0 -Sc 0 5 0 0 $s_2$ -Sc 0 6 0 0 $s_2$ -Sc 0 7 0 0 $s_2$ -Sc 0 8 0 0 $s_2$ -Sp 0.5 -SFC

**$\mathcal{M}_3$ (neutral demography)**

msms 80 1 -t 10 -I 2 40 40 -n 1 1 -n 2 1 -m 1 2 $M$ -m 2 1 $M$ -em 0.5 1 2 0 -em 0.5 2 1 0 -eg 4 1 2 -ej 5 1 2 -N 10000

**$\mathcal{M}_3$ (selection addendum)**

-SI $T_S$ 2 $f_1$ $f_2$ -Sc 0 1 $s_1$ 0 0 -Sc 0 2 0 0 $s_2$ -Sp 0.5 -SFC

**$\mathcal{M}_4$ (neutral demography)**

msms 160 1 -t 10 -I 4 40 40 40 40 -n 1 0.01 -n 2 0.1 -n 3 5.0 -n 4 0.1 -m 1 2 2 -m 2 1 2 -m 2 3 2 -m 3 2 2 -m 3 4 2 -m 4 3 2 -m 4 2 $M$ -m 2 4 $M$ -em 1.0 2 4 0 -em 1.0 4 2 0 -en 1.5 3 1 -g 3 2.5 -ej 1 1 2 -ej 1.5 2 3 -ej 4 4 3 -N 10000

**$\mathcal{M}_4$ (selection addendum)**

-SI $T_S$ 4 $f_1$ $f_2$ $f_3$ $f_4$ -Sc 0 1 0 0 0 -Sc 0 2 $s_2$ 0 0 -Sc 0 3 0 0 0 -Sc 0 4 0 0 $s_4$ -Sp 0.5 -SFC

#### S3. LSD command lines for calculating summary statistics for models $\mathcal{M}_1$, $\mathcal{M}_2$, $\mathcal{M}_3$ and $\mathcal{M}_4$

$\mathcal{M}_1$

python3 lsd_hi.py ${*msms_output*} -d 40 -d 40 -l 5000 -f ABC

$\mathcal{M}_2$

python3 lsd_hi.py ${*msms_output*} -d 40 -d 40 -d 40 -d 40 -d 40 -d 40 -l 5000 -f ABC

$\mathcal{M}_3$

python3 lsd_hi.py ${*msms_output*} -d 40 -d 40 -l 5000 -f ABC

$\mathcal{M}_4$

python3 lsd_hi.py ${*msms_output*} -d 40 -d 40 -d 40 -d 40 -l 5000 -f ABC

#### S4. Estimating neutral demographic parameters $\hat{\boldsymbol{\theta}}$

In LSD, estimation of model parameters is performed in windows. That is, the genome (full dataset) is partitioned into windows (subsets) on which ABC parameter estimation is performed. We posit that for those windows evolving exclusively under neutrality, the underlying model parameters should be shared and be reflective of the full neutral genome partition. In such a case, we aim to estimate the posterior density given the full neutral region partition (dataset) by combining the neutral window (subset) posterior samples. Several methods have been developed that attempt to do this. Here, we compare the methods of i) average of subposterior samples, ii) consensus Monte Carlo, assuming independence of individual model parameters (Scott et al., 2016), iii) consensus Monte Carlo, assuming covariance among model parameters (Scott et al., 2016), iv) semi-parametric density product estimator (Neiswanger, Wang, & Xing, 2013), v) product of densities, vi) independent replicates (Thalmann et al., 2011) and vii) independent replicates MCMC (Thalmann et al., 2011). The first four are implemented in the R package *parallelMCMCcombine* (Miroshnikov & Conlon, 2014) and the last two in *ABCtoolbox* (Wegmann, Leuenberger, Neuenschwander, & Excoffier, 2010). Using a simulated pseudo-genome under model $\mathcal{M}_2$, and given 500 neutral 5kb windows, we show the method of product of densities to be the most consistently accurate across model parameters (truth values indicated in dashed black lines; Figure S2), in addition to being by a considerable margin the fastest to calculate. However, given that this method does not return a true posterior, we opt to take the mode of this distribution as the single point estimator $\hat{\theta}$ in our current LSD implementation.

#### S5. Robustness of method to misspecification of neutral windows

Given that LSD relies on a two-step process of first acquiring neutral model parameter estimates and then estimating model parameters in a genome scan, it is pertinent to assess the robustness of LSD to mis-specification of neutral windows in the estimation of neutral model parameters. Here, for each model-selection regime combination, we simulate 1000 neutral and 500 selected 5kb loci. From this we construct a 1000 loci (5Mb) pseudo-genome comprising a fraction $f$ of selected loci and a fraction 1-$f$ of neutral loci, with $0.0 \leq f \leq 0.2$, by drawing loci from the larger simulated sets. Pseudo-genomes were generated under all four demographic models ($\boldsymbol{\mathcal{M}_1}, \dots, \boldsymbol{\mathcal{M}_4}$) under neutral migration rates $M_{12}, M_{21} = 5$ and under medium and strong selection coefficients ($s_1 = s_2 = s = 0.01, 0.1$; $T_S = 40{,}000$ generations ago). We then estimate the neutral $\widehat{M}_{12}$, $\widehat{M}_{21}$ parameters utilising the 1000 loci pseudo-genome, and calculate the two-dimensional Euclidean distance of the $\widehat{M}_{12}$, $\widehat{M}_{21}$ reciprocal migration parameters from the truth (**Figure S3**). Reciprocal migration parameters have same prior range and scale. We observe in **Figure S3** that under most models ($\boldsymbol{\mathcal{M}_1}$, $\boldsymbol{\mathcal{M}_2}$ and $\boldsymbol{\mathcal{M}_3}$), there is minimal increase in error (distance from the truth) with increasing fraction of selected loci included in the neutral set. This may be attributed to our method of estimating $\widehat{\boldsymbol{\theta}}$ via the product of densities, which amplifies the signal (density) according to majority rule. Models $\boldsymbol{\mathcal{M}_1}$ and $\boldsymbol{\mathcal{M}_3}$ show relatively small absolute errors across that range ($\leq$ 0.5) while $\boldsymbol{\mathcal{M}_2}$ shows circa an order of magnitude difference from the truth. Unlike the other models, model $\boldsymbol{\mathcal{M}_4}$ shows a positive cline of increasing error with increasing fraction of selected sites, with 0.2 error at 0% selected sites increasing to an error of 2.5 at 20%.

We note however that the errors under models $\boldsymbol{\mathcal{M}_2}$ and $\boldsymbol{\mathcal{M}_4}$ (at high fractions of selected windows) do not necessarily mean that LSD's power to detect selection under these models is reduced. This is because what dictates the performance of LSD are not the absolute values of the parameter estimates but rather the relative distance between the neutral parameter estimates $\widehat{\boldsymbol{\theta}}$ and the window posteriors $\pi_l(\boldsymbol{\theta})$, specifically that of windows under selection. To demonstrate this, we calculate the AUC of pseudo-genomes comprising 1000 neutral and 50 selected loci, under values of $\widehat{\boldsymbol{\theta}}$ that were estimated from a neutral set comprising a fraction $f$ of selected loci and a fraction 1-$f$ of neutral loci, with $0.0 \leq f \leq 0.2$, by drawing loci from the larger simulated sets as done previously; for the same set of demographic models and selection regimes. Here, the selected loci comprising the pseudo-genome under the selection scan are under the same selection regime as those included as mis-specifications in the neutral set. We show that the power to detect selection under LSD is very robust to mis-specification of the neutral set, with negligible slopes for model $\boldsymbol{\mathcal{M}_1}, \boldsymbol{\mathcal{M}_2}$ and $\boldsymbol{\mathcal{M}_3}$ across the range of fractions (**Figure 4**). For model $\boldsymbol{\mathcal{M}_4}$, the slope is minimal up until 13% mis-specification, after which we observed a steep negative slope towards an AUC of 0.5 at 17% mis-specification. While the degree of robustness appears to depend on the complexity of the demographic model, such that complex demographies are less forgiving towards mis-specification of the neutral set, LSD appear to tolerate levels of mis-specifications beyond what is generally expected in the genomes of most biological systems, where elevated proportions of non-neutral sites generally imply selection coefficients of small size on many loci, such as in polygenic adaptation or background selection.

#### S6. msms and LSD command lines for generating simulated data and simulated summary statistics, for models $\mathcal{M}_1$ and $\mathcal{M}_2$ for the case study

For msms, $\theta = 4 \times N_{msms} \times mutation\,rate \times window\,size = 4 \times 10000 \times 1.7 \text{ x } 10^{-8} \times 10000 = 6.8$; $N$ is in fraction of $N_{msms}$ (= 10000), and the mutation rate ($1.7 \text{ x } 10^{-8}$) is the same as that used in Tavares et al. 2018.

For LSD-Hi, we simulate pooled data (-p), sequencing errors (-i --error_method 4 --error_rate 0.001) and the same filtering regime as in Tavares et al. 2018 (--minallelecount 2 --mindepth 15 --maxdepth 500). We further replicated the coverage distribution of the observed data by drawing samples from a parametric (negative binomial) distribution fitted to the empirical coverage distribution (--sampler nbinom -c *dist_file*)

##### $\mathcal{M}_1$

msms 200 1 -t $\theta$ -I 2 100 100 -n 1 $N_{YP1}$ -n 2 $N_{MP2}$ -m 1 2 $M_{YP1\_MP2}$ -m 2 1 $M_{MP2\_YP1}$

python3 lsd_hi.py ${*msms_output*} -d 100 -d 100 -l 10000 -p -i --error_method 4 --error_rate 0.001 --minallelecount 2 --mindepth 15 --maxdepth 500 --sampler nbinom -c Amajus_coverage_simpleModel_nbinomDist.txt -f ABC

##### $\mathcal{M}_2$

msms 598 1 -t $\theta$ -I 8 84 100 100 0 0 100 98 116 -n 1 $N_{CAM}$-n 2 $N_{ML}$ -n 3 $N_{YP1}$ -n 4 $N_Y$ -n 5 $N_R$ -n 6 $N_{MP2}$ -n 7 $N_{CHI}$ -n 8 $N_{CIN}$ -m 1 4 $M_{ic}$ -m 2 4 $M_{ic}$ -m 3 4 $M_{ic}$ -m 8 5 $M_{ic}$ -m 7 5 $M_{ic}$ -m 6 5 $M_{ic}$ -m 4 5 $M_{YR}$ -m 5 4 $M_{RY}$

python3 lsd_hi.py ${*msms_output*} -d 84 -d 100 -d 100 -d 100 -d 98 -d 116 -l 10000 -p -i --error_method 4 --error_rate 0.001 --minallelecount 2 --mindepth 15 --maxdepth 500 --sampler nbinom -c Amajus_coverage_6popsIMmodel_nbinomDist.txt -f ABC

#### S7. LSD command lines for calculating summary statistics from observed data, for models $\mathcal{M}_1$ and $\mathcal{M}_2$ of the case study

We calculate summary statistics for the observed data (neutral regions and chromosome 6) considering sequences are pooled (–pooled), for a 10kb window size (in addition to 1kb step-size for the scan) (--windowSize 10000 (--windowStep 1000)) and the same filtering regime as in the simulated data (--minallelecount 2 --mindepth 15 --maxdepth 500).

**$\mathcal{M}_1$ (chromosome 6)**

```
python lsd_high_sumstats_calculator_OBS.py Chr6_simpleModel_filelist.txt -d 100 -d 100 -q 0 -m 2 -o Chr6_2pops_simpleModel -f ABC -r single --startPos 1 --endPos 55771888 --mindepth 15 --maxdepth 500 --windowSize 10000 --windowStep 1000 –pooled
```

**$\mathcal{M}_1$ (neutral regions)**

```
python lsd_high_sumstats_calculator_OBS.py Chr6_neutral_simpleModel_filelist.txt -d 100 -d 100 -q 0 -m 2 -o neutral_Chr6_2pops_simpleModel -f ABC -r single --startPos 1 --endPos 55771888 --mindepth 15 --maxdepth 500 --windowSize 10000 --pooled
```

**$\mathcal{M}_2$ (neutral regions)**

```
python lsd_high_sumstats_calculator_OBS.py Chr6_6popIMmodel_filelist.txt -d 84 -d 100 -d 100 -d 100 -d 98 -d 116 -q 0 -m 2 -o Chr6_6popIM -f ABC -r single --startPos 1 --endPos 55771888 --mindepth 15 --maxdepth 500 --windowSize 10000 --windowStep 1000 –pooled
```

**$\mathcal{M}_2$ (chromosome 6)**

```
python lsd_high_sumstats_calculator_OBS.py Chr6_neutral_6popIMmodel_filelist.txt -d 84 -d 100 -d 100 -d 100 -d 98 -d 116 -q 0 -m 2 -o neutral_Chr6_6popIMmodel -f ABC -r single --startPos 1 --endPos 55771888 --mindepth 15 --maxdepth 500 --windowSize 10000 --pooled
```

### SUPPLEMENTARY FIGURES

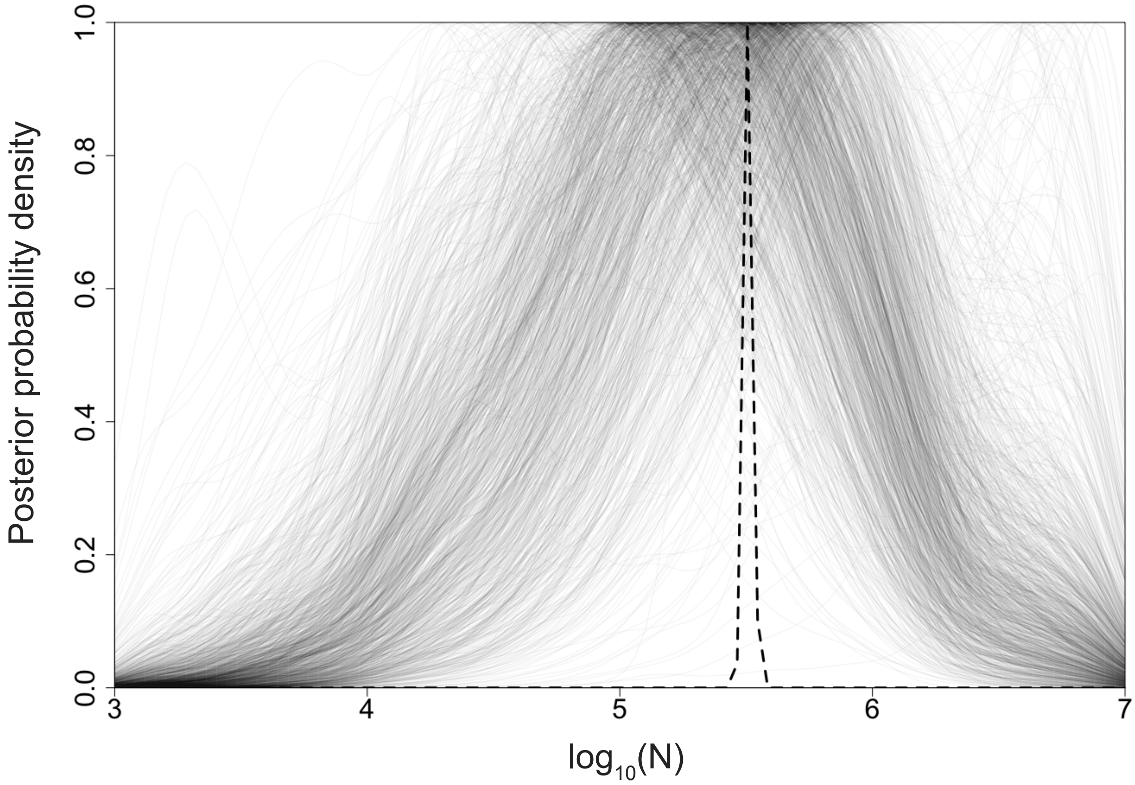


**Figure S1.** Estimation of $\boldsymbol{\hat{\theta}}$ as the mode of the product (black dashed line) of the local posterior densities of 1000 putatively neutral (5kb) windows (grey solid lines).

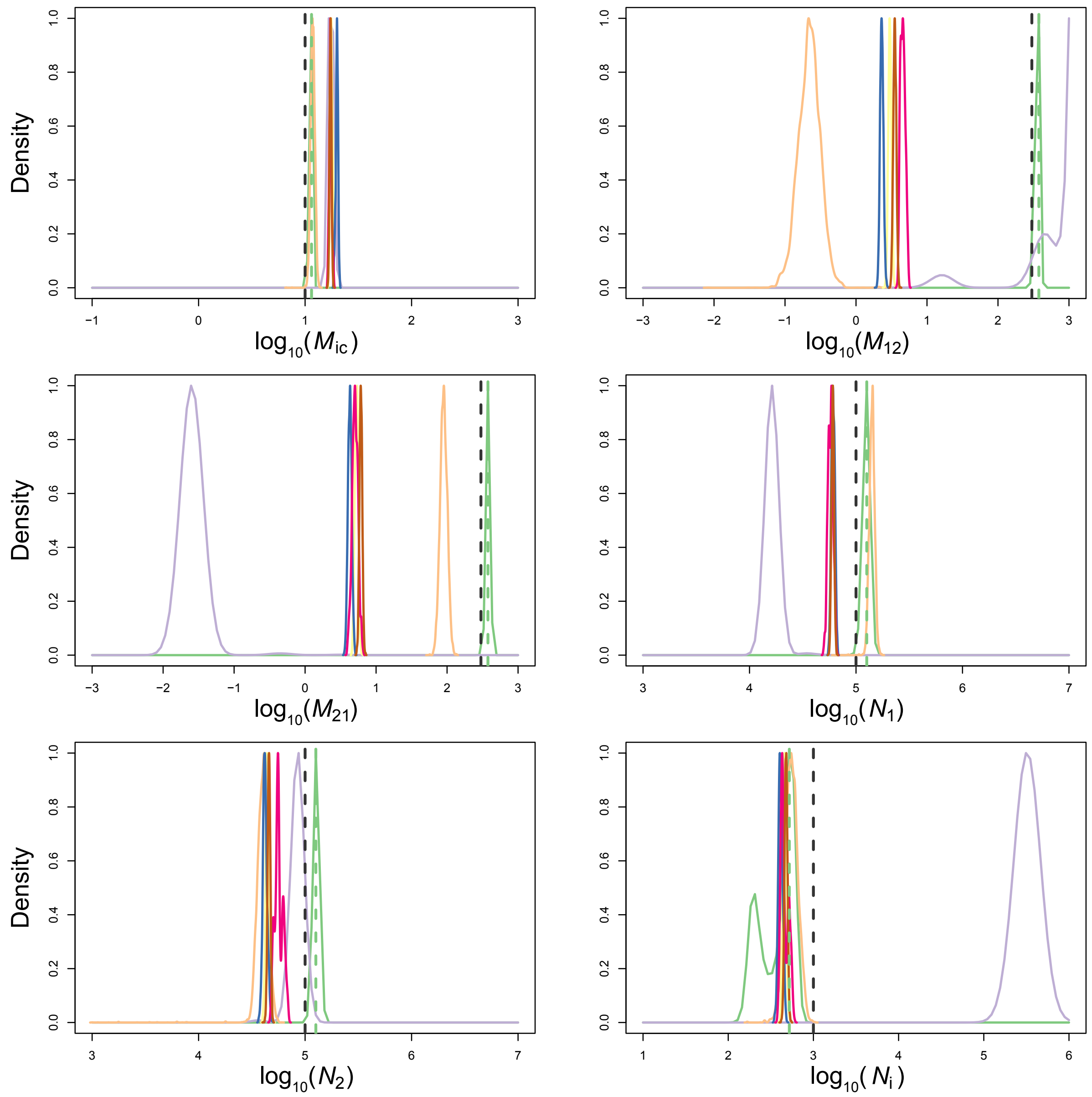


**Figure S2.** Comparison of different methods for estimating the posterior density given the full neutral region partition by combining the neutral window (subset) posterior samples. This comparison was performed under a simulated pseudo-genome (model $\mathcal{M}_2$) comprised of 500 neutral (5kb) windows. Seven methods are compared: i) average of subposterior samples (pink), ii) consensus Monte Carlo, assuming independence of individual model parameters (blue), iii) consensus Monte Carlo, assuming covariance among model parameters (yellow), iv) semi-parametric density product estimator (brown) v) product of densities (green), vi) independent replicates (purple) and vii) independent replicates MCMC (gold). The mode of product of densities (green, dashed) and the ground truth (black, dashed) are also shown. $x$-axes ranges reflect model parameter prior ranges.

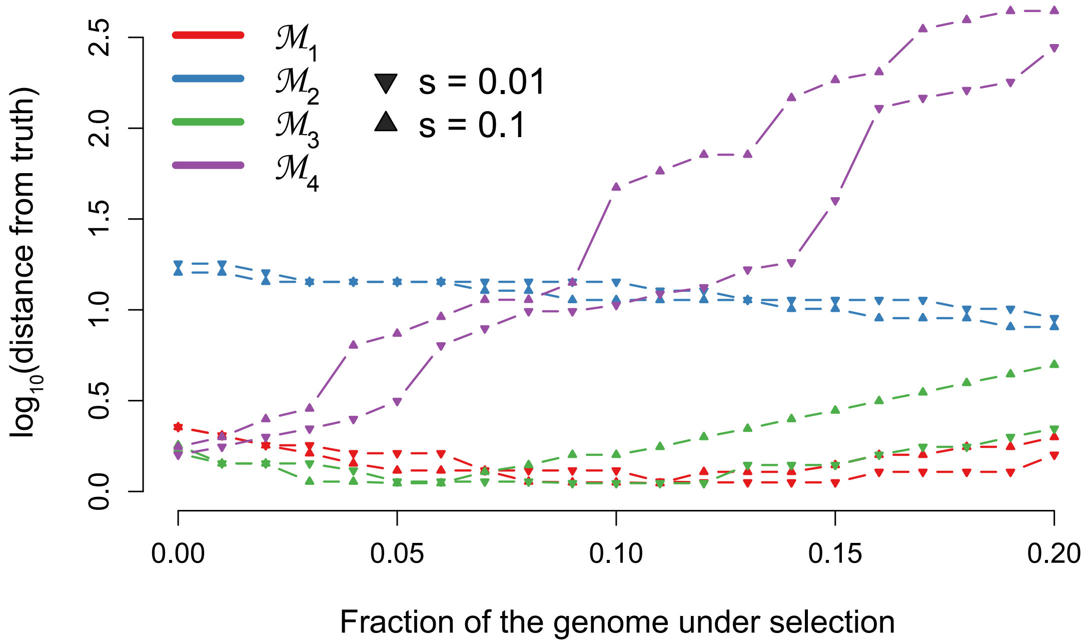


**Figure S3.** Effect of increasing fraction of mis-specified (aka selected) windows among the neutral set on error, for all 4 models and intermediate and high selection coefficients ($s_1 = s_2 = s$ = 0.01, 0.1; $T_S$ = 40,000) under neutral migration rates $M_{12}, M_{21} = 5$. Here the neutral set of 1000 loci comprise a fraction *f* of selected loci and a fraction 1-*f* neutral loci, with $0.0 \leq f \leq 0.2$. The metric of error (*y*-axis) is defined as the $\log_{10}$ 2-dimensional Euclidean distance of the reciprocal migration parameters $\widehat{M}_{12}, \widehat{M}_{21}$ as estimated from combining all windows via their product of densities (the neutral estimate) from the true (simulated) neutral migration rates.

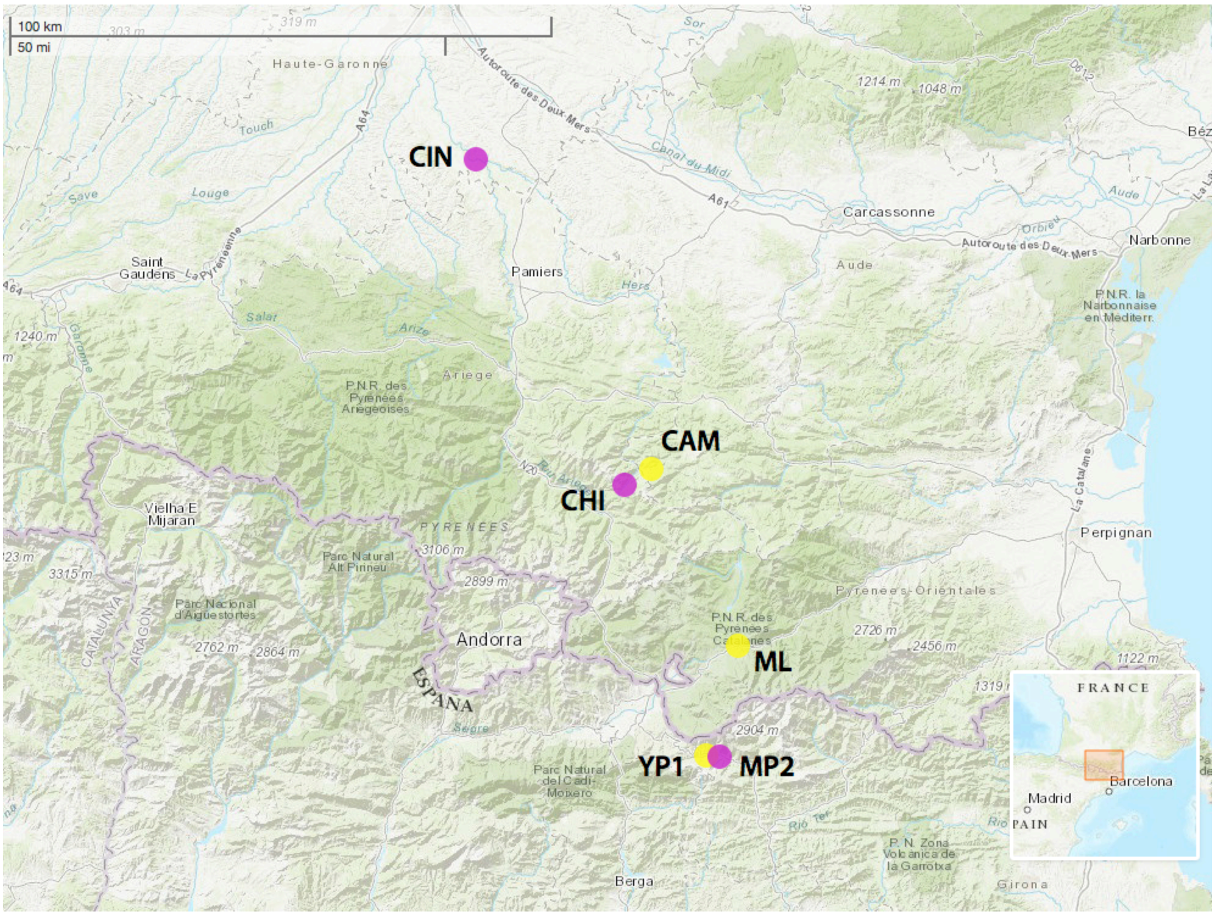


**Figure S4.** Map of the *A.m.pseudomajus* (magenta) and *A.m.striatum* (yellow) populations used in models $\mathcal{M}_1$ and $\mathcal{M}_2$ of the case study. These populations were sequenced via pool-seq in a previous study (Tavares et al., 2018).

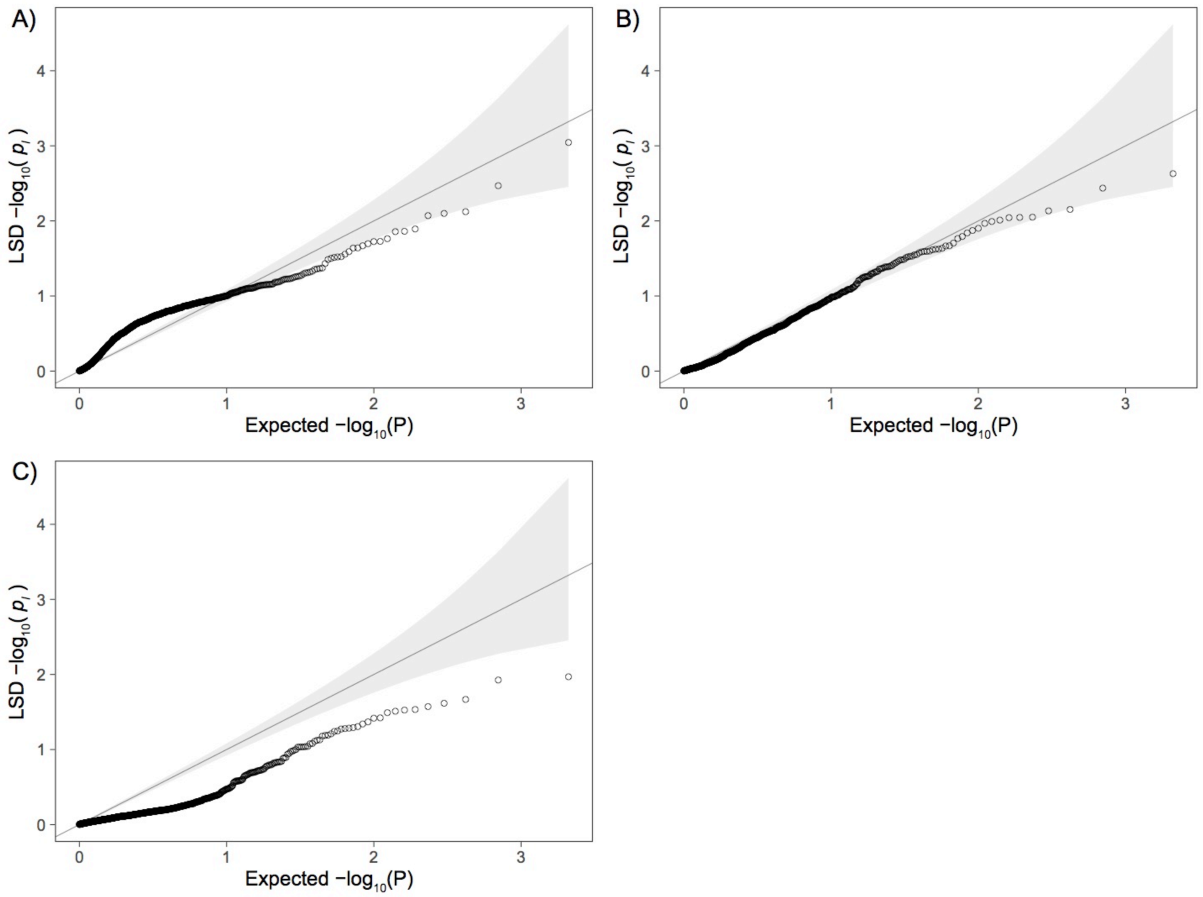


**Figure S5.** Quantile-quantile (Q-Q) plots comparing LSD's $p_l$ probability distribution against that of expectation, evaluated on neutral-exclusive (1000 loci) pseudo-genomes for model $\boldsymbol{\mathcal{M}_1}$, under neutral migration rates A) $M = 0.5$, B) $M = 5$, and C) $M = 50$. Grey shaded areas represent expected 95% confidence intervals.

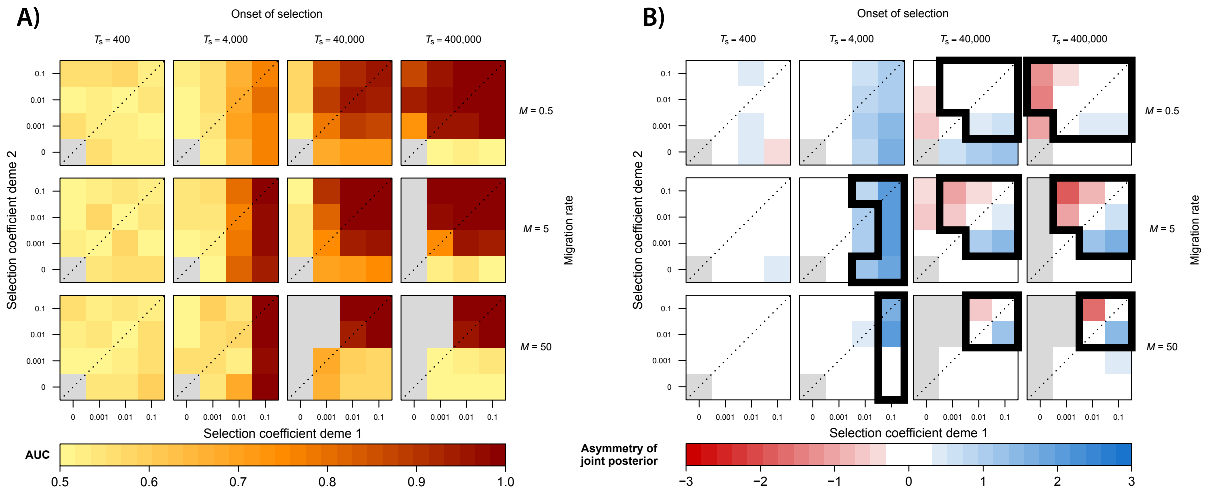


**Figure S6.** Simulation results for the 2-deme IM model (model $\mathcal{M}_1$; *de-novo* case) showing the effect of migration rate, time of onset of selection and deme-specific selection coefficients on A) LSD diagnostic performance (AUC) and B) LSD inferred (a)symmetry of selection. Each coloured cell represents a pseudo-genome simulated under a specific selection regime. Grey cells indicate selection regimes where the derived allele is always lost. A) The cell colours reflect the AUC value of the LSD scan calculated by the correct discrimination of 1000 neutral loci and 50 selected loci in the 1050 loci simulated pseudo-genomes. An AUC value of 0.5 reflects random assignment while that of 1 reflects perfect classification (i.e. TPR=1, FPR=0). B) The cell colours reflect the (a)symmetry values inferred by LSD, where a value of 0 reflects perfect symmetry of the joint posterior while values divergent from this reflect asymmetry. Cells surrounded by thick lines indicate the values of (a)symmetry for regimes expected to generate meaningful signal (AUC > 0.8 in Figure S6A).

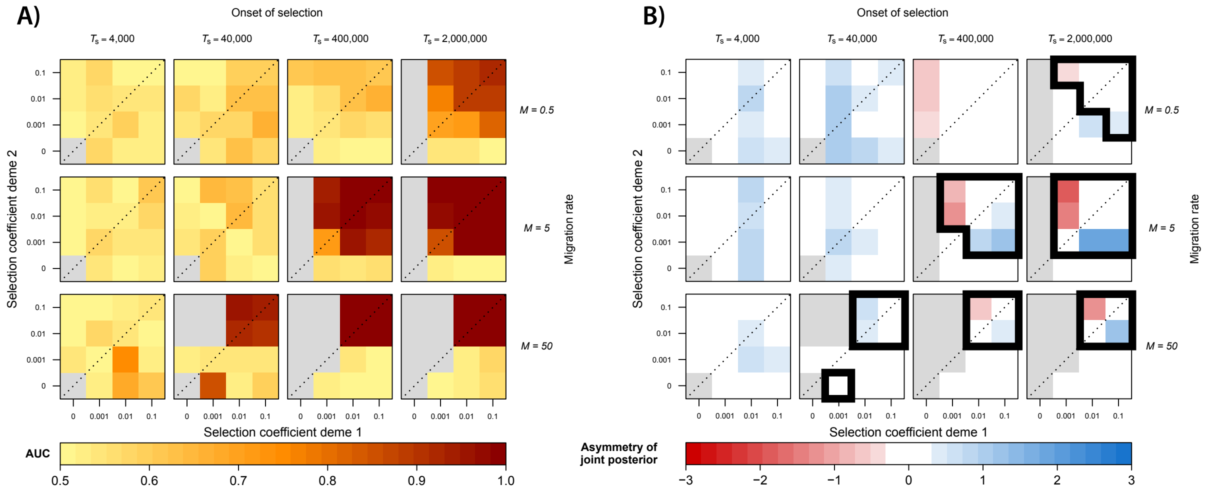


**Figure S7.** Simulation results for the 6-deme IM model (model $\mathcal{M}_2$; standing variation case) showing the effect of migration rate, time of onset of selection and deme-specific selection coefficients on A) LSD diagnostic performance (AUC) and B) LSD inferred (a)symmetry of selection. Each coloured cell represents a pseudo-genome simulated under a specific selection regime. Grey cells indicate selection regimes where the derived allele is always lost. A) The cell colours reflect the AUC value of the LSD scan calculated by the correct discrimination of 1000 neutral loci and 50 selected loci in the 1050 loci simulated pseudo-genomes. An AUC value of 0.5 reflects random assignment while that of 1 reflects perfect classification (i.e. TPR=1, FPR=0). B) The cell colours reflect the (a)symmetry values inferred by LSD, where a value of 0 reflects perfect symmetry of the joint posterior while values divergent from this reflect asymmetry. Cells surrounded by thick lines indicate the values of (a)symmetry for regimes expected to generate meaningful signal (AUC > 0.8 in Figure S7A).

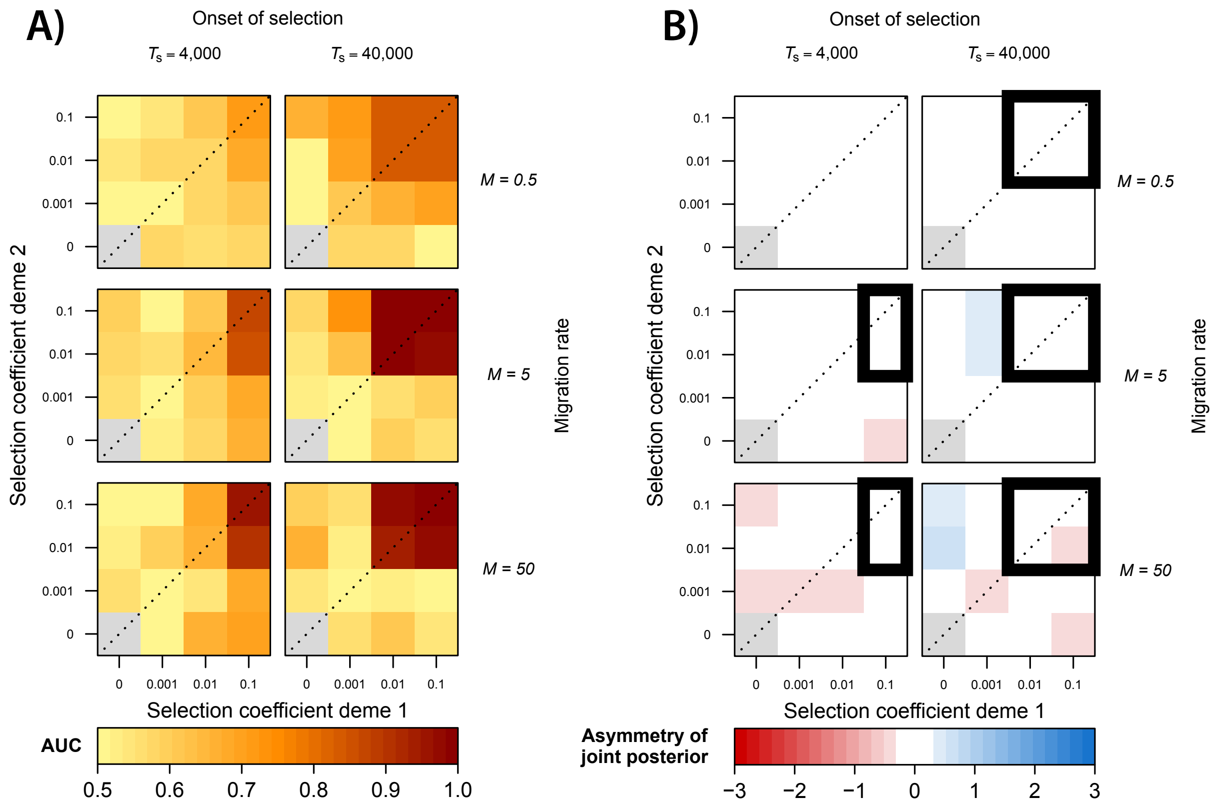


**Figure S8.** Simulation results for the 4-deme hierarchical divergence model (model $\mathcal{M}_4$; standing variation case) showing the effect of migration rate, time of onset of selection and deme-specific selection coefficients on A) LSD diagnostic performance (AUC) and B) LSD inferred (a)symmetry of selection. Each coloured cell represents a pseudo-genome simulated under a specific selection regime. Grey cells indicate selection regimes where the derived allele is always lost. A) The cell colours reflect the AUC value of the LSD scan calculated by the correct discrimination of 1000 neutral loci and 50 selected loci in the 1050 loci simulated pseudo-genomes. An AUC value of 0.5 reflects random assignment while that of 1 reflects perfect classification (i.e. TPR=1, FPR=0). B) The cell colours reflect the (a)symmetry values inferred by LSD, where a value of 0 reflects perfect symmetry of the joint posterior while values divergent from this reflect asymmetry. Cells surrounded by thick lines indicate the values of (a)symmetry for regimes expected to generate meaningful signal (AUC > 0.8 in Figure S8A).

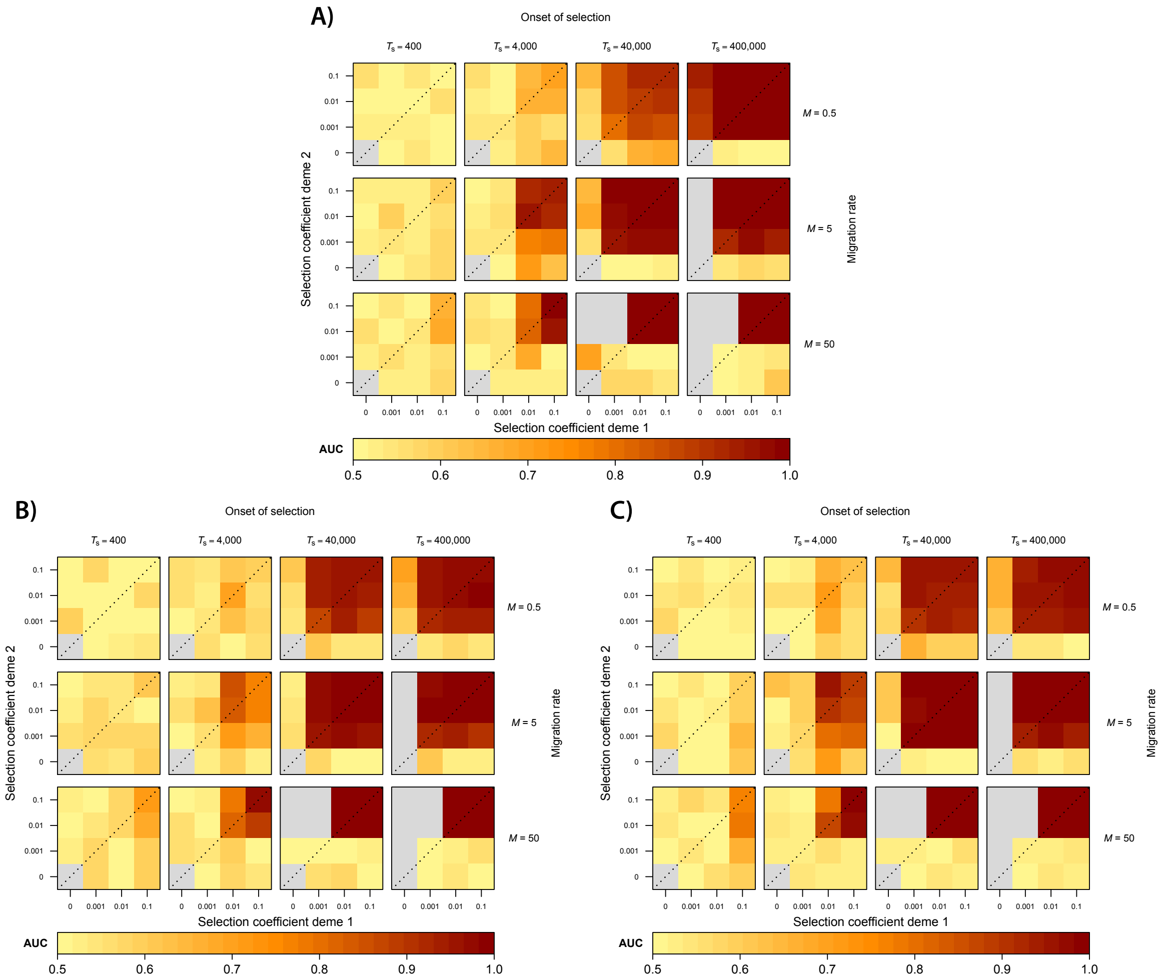


**Figure S9.** Comparison of power to detect selection (AUC) between (A) LSD, (B) pcadapt and (C) OutFLANK under model $\boldsymbol{\mathcal{M}_1}$. For LSD, two parametrisations are performed: i) assuming non-$M$ demographic parameters and neutral $\widehat{M}$ to be fixed to the true values (LSD FIXED, Figure 4) and ii) allowing non-$M$ demographic parameters to be drawn from a large prior range and using neutral $\widehat{M}$ estimated from the pseudo-genomes (LSD FREE, Figure S9A). Grey cells indicate selection regimes where the derived allele is always lost.

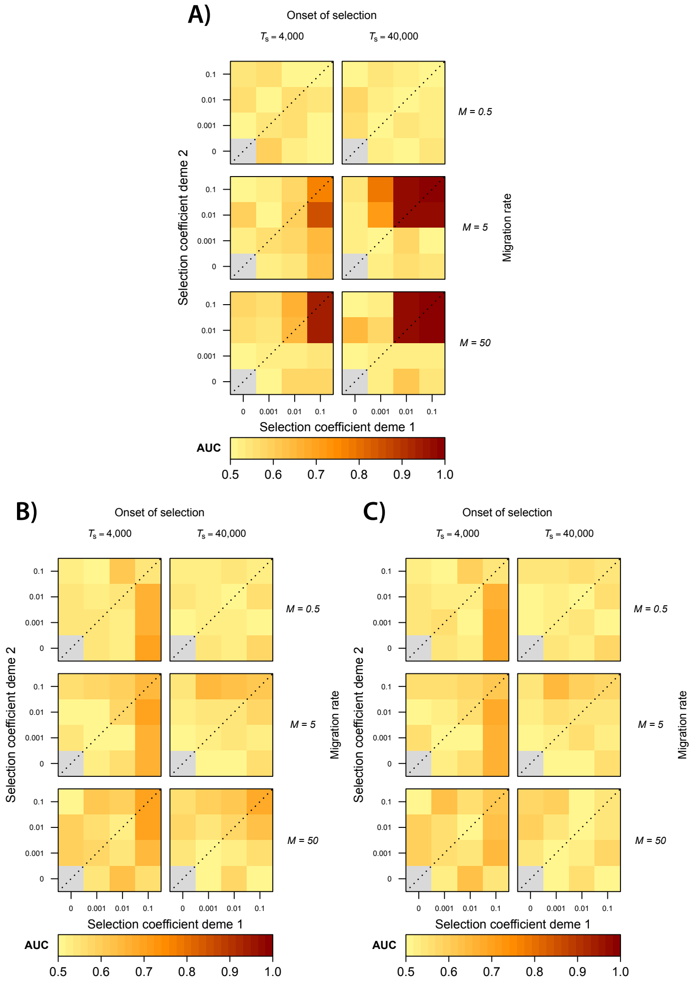


**Figure S10.** Comparison of power to detect selection (AUC) between (A) LSD, (B) pcadapt and (C) OutFLANK under model $\mathcal{M}_4$. For LSD, two parametrisations are performed: i) assuming non-$M$ demographic parameters and neutral $\widehat{M}$ to be fixed to the true values (LSD FIXED, Figure S8A) and ii) allowing non-$M$ demographic parameters to be drawn from a prior range and using neutral $\widehat{M}$ estimated from the pseudo-genomes (LSD FREE, Figure S10A). Grey cells indicate selection regimes where the derived allele is always lost.

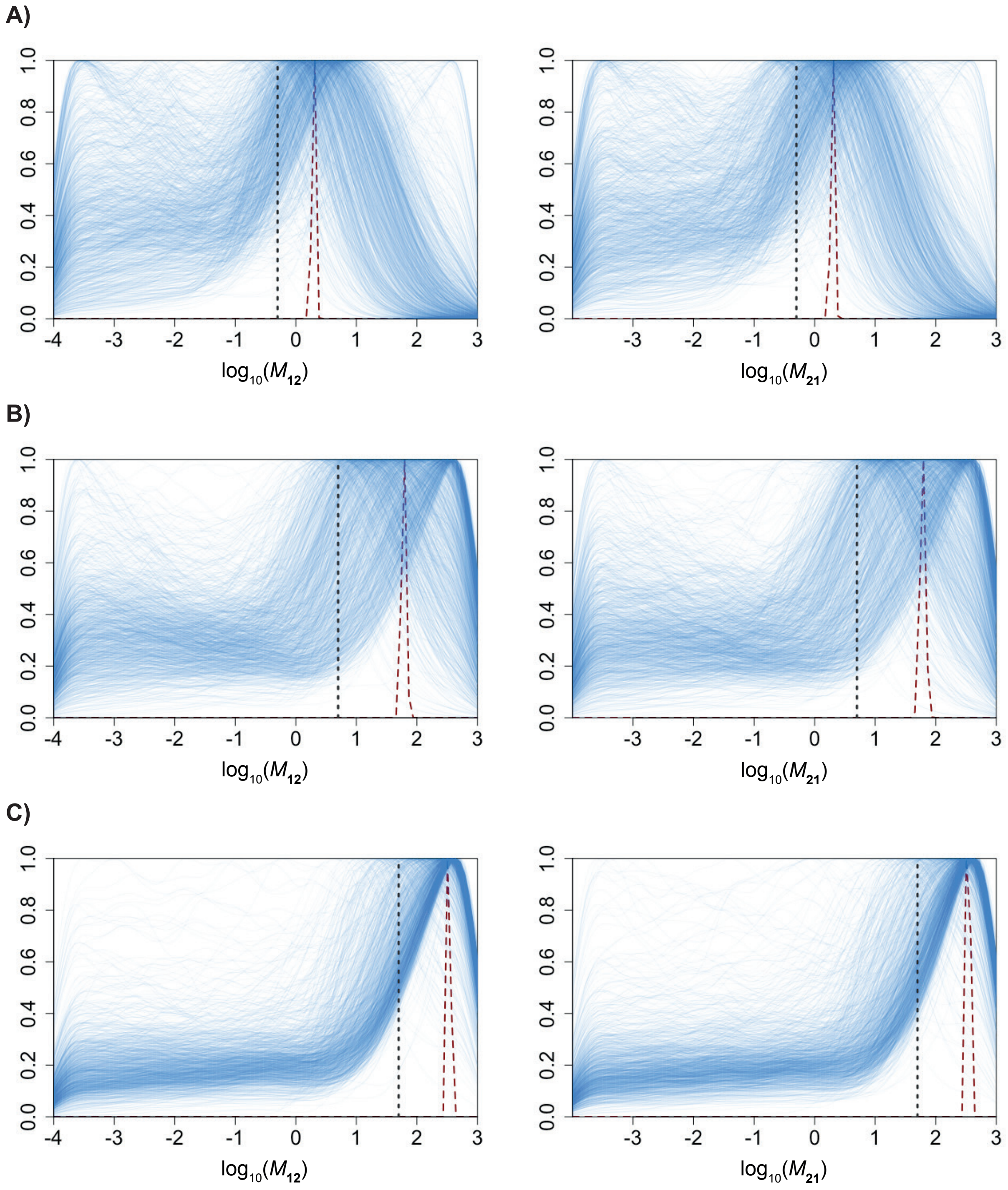


**Figure S11.** Estimation of $\widehat{M}$ (red dashed lines) from the local posteriors of 1050 5kb windows (blue solid lines) for model $\boldsymbol{\mathcal{M}_1}$ “free” parameterisation (LSD FREE), under neutral $M_{12} = M_{21} = M = 0.5$ (A) 5 (B) and 50 (C) 50. For added realism, the neutral set comprised 1,000 neutral and 50 weakly selected ($s_1, s_2 = 0.001$; $T_S = 4{,}000$) aka mis-specified loci. The true $M$ values are indicated by the black dashed lines. $M_{12}, M_{21}$ are scaled to a global $N_E$ (=10,000). $x$-axes ranges reflect prior ranges.

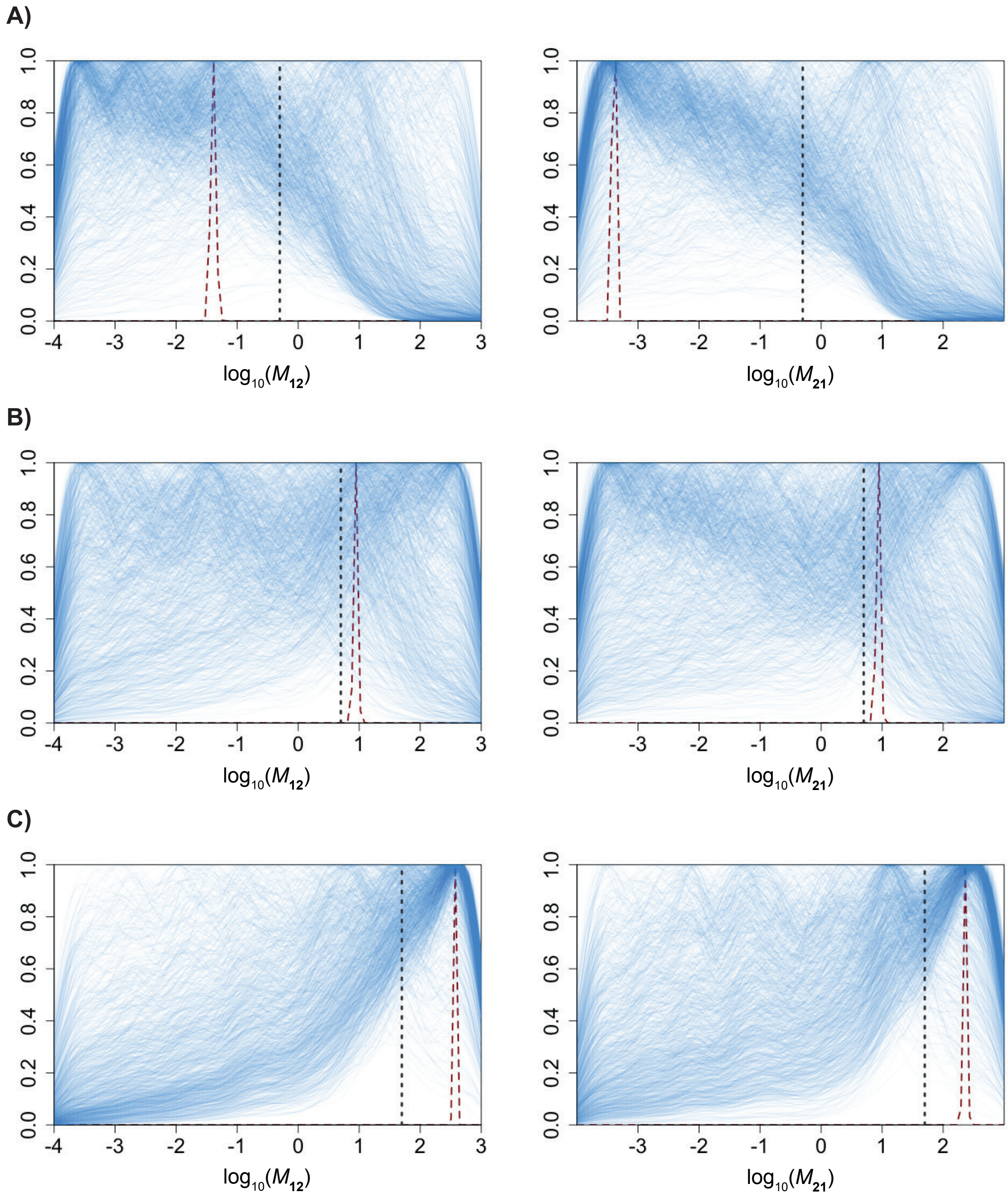


**Figure S12.** Estimation of $\widehat{M}$ (red dashed lines) from the local posteriors of 1050 5kb windows (blue solid lines) for model $\boldsymbol{\mathcal{M}_4}$ “free” parameterisation (LSD FREE), under neutral $M_{12} = M_{21} = M = 0.5$ (A) 5 (B) and 50 (C) 50. For added realism, the neutral set comprised of 1,000 neutral and 50 weakly selected ($s_{1,}\ s_2 = 0.001$; $T_S = 4{,}000$) aka mis-specified loci. The true $M$ values are indicated by the black dashed lines. $M_{12}, M_{21}$ are scaled to a global $N_E$ (=10,000). $x$-axes ranges reflect prior ranges.

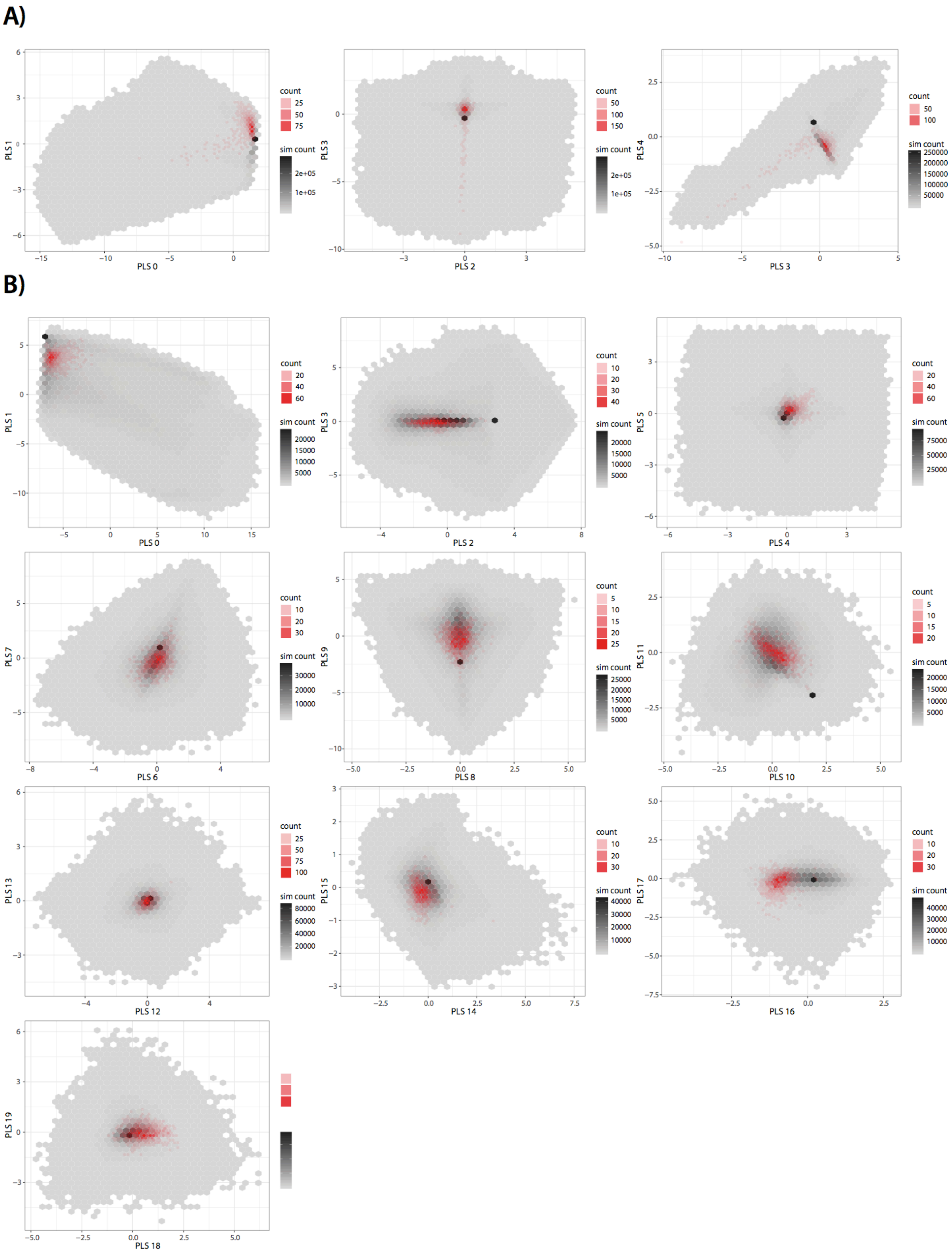


**Figure S13.** In an ABC implementation of LSD, models should be validated by demonstrating that simulations capture the range of, as well as be centred around, the observed data to be maximally informative for parameter estimation. Here we show high density scatter plots with binning (hexbin plots) of observed (neutral) summary statistics (red)

overlaid on simulated summary statistics (grey), for the *A. majus* LSD scans for A) model $\mathcal{M}_1$ (decomposed into 5 PLS components) and B) model $\mathcal{M}_2$ (decomposed into 20 PLS components).

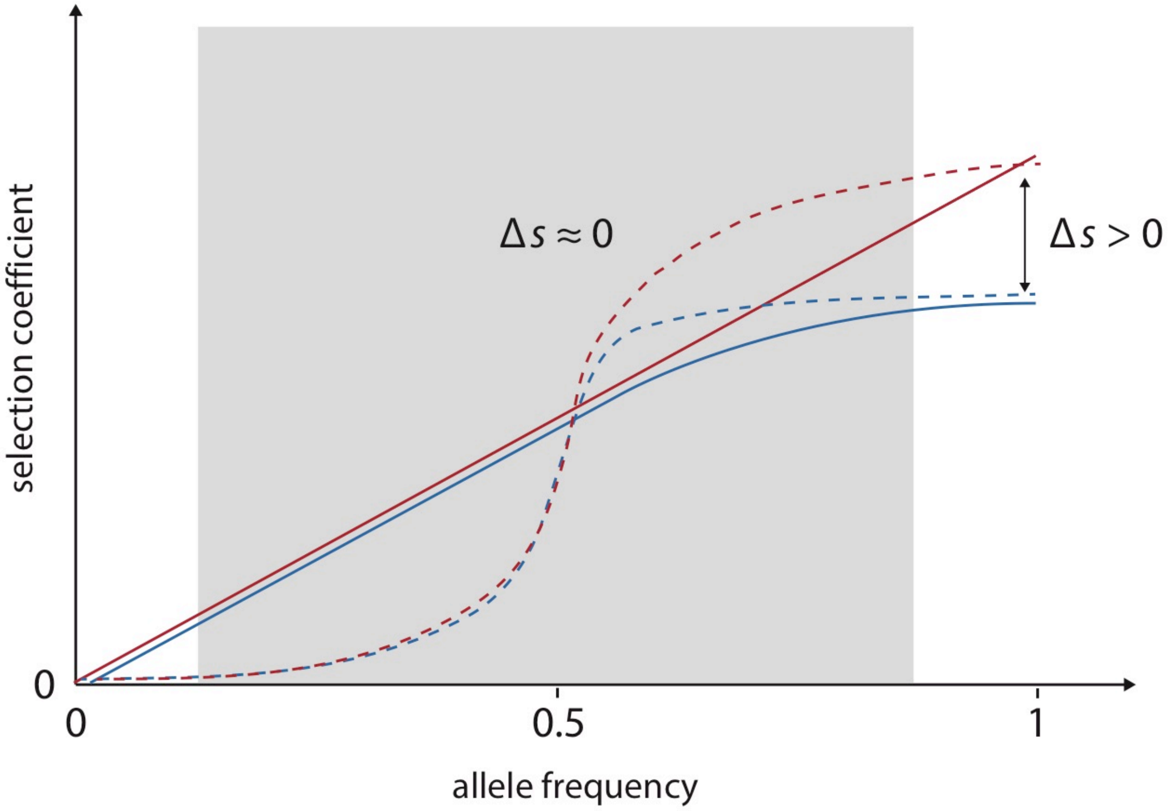


**Figure S14.** Selection on alternate alleles (blue and red) with differing positive frequency-dependent curves (regimes) could underlie the difference in inferred (a)symmetry between models $\mathcal{M}_1$ (population pair near contact zone, $\Delta s \approx 0$) and $\mathcal{M}_2$ (global differences between subspecies, $\Delta s > 0$) for the ROS and EL loci in the case study. Examples are shown for pseudo-linear (solid lines) and logistic (dashed lines) curve pairs. The grey-shaded interval represents allele frequencies that characterise the contact zone.

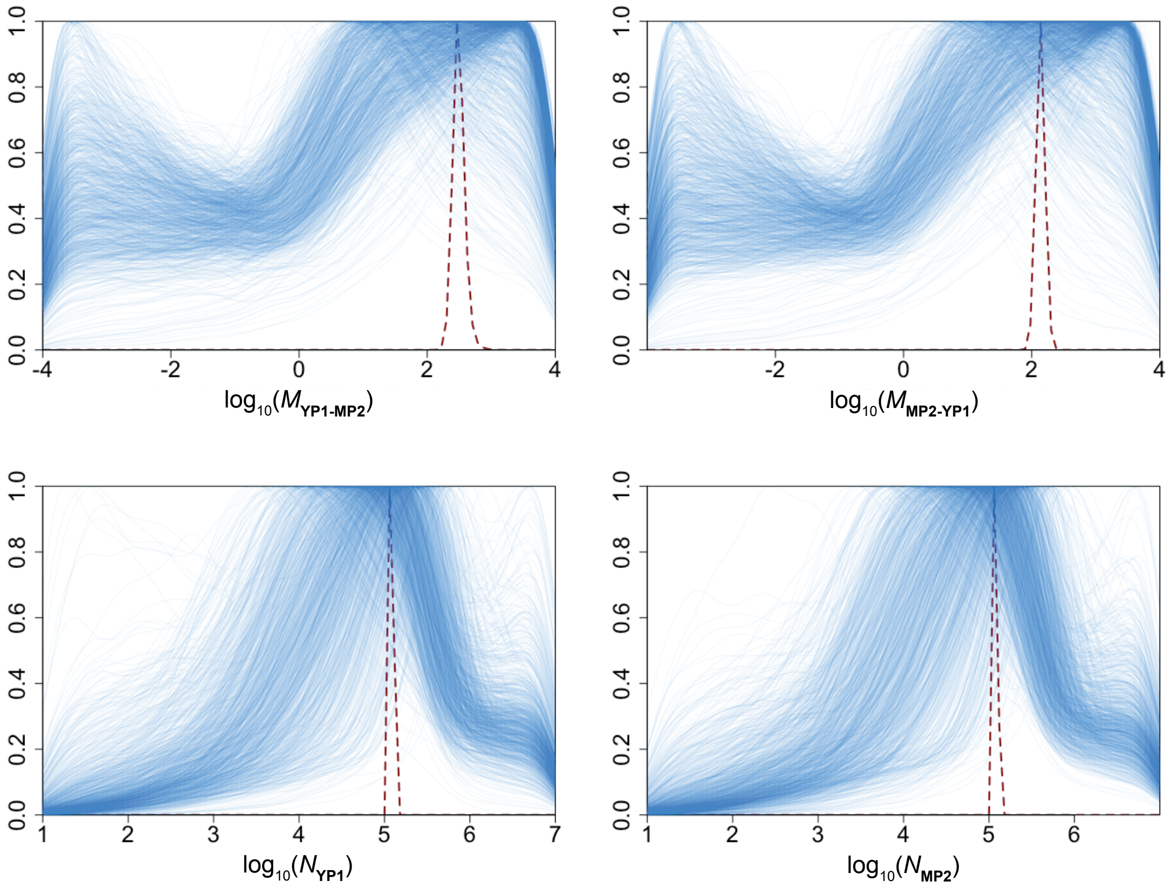


**Figure S15.** Estimation of $\widehat{\boldsymbol{\theta}}$ (red dashed lines) from the posteriors of 1210 10kb putatively neutral windows (blue solid lines) for model $\boldsymbol{\mathcal{M}_1}$ in the *A.majus* case study. $M_{YP1-MP2}, M_{MP2-YP1}$ are scaled to a global $N_E$ (=10,000). $x$-axes ranges reflect prior ranges.

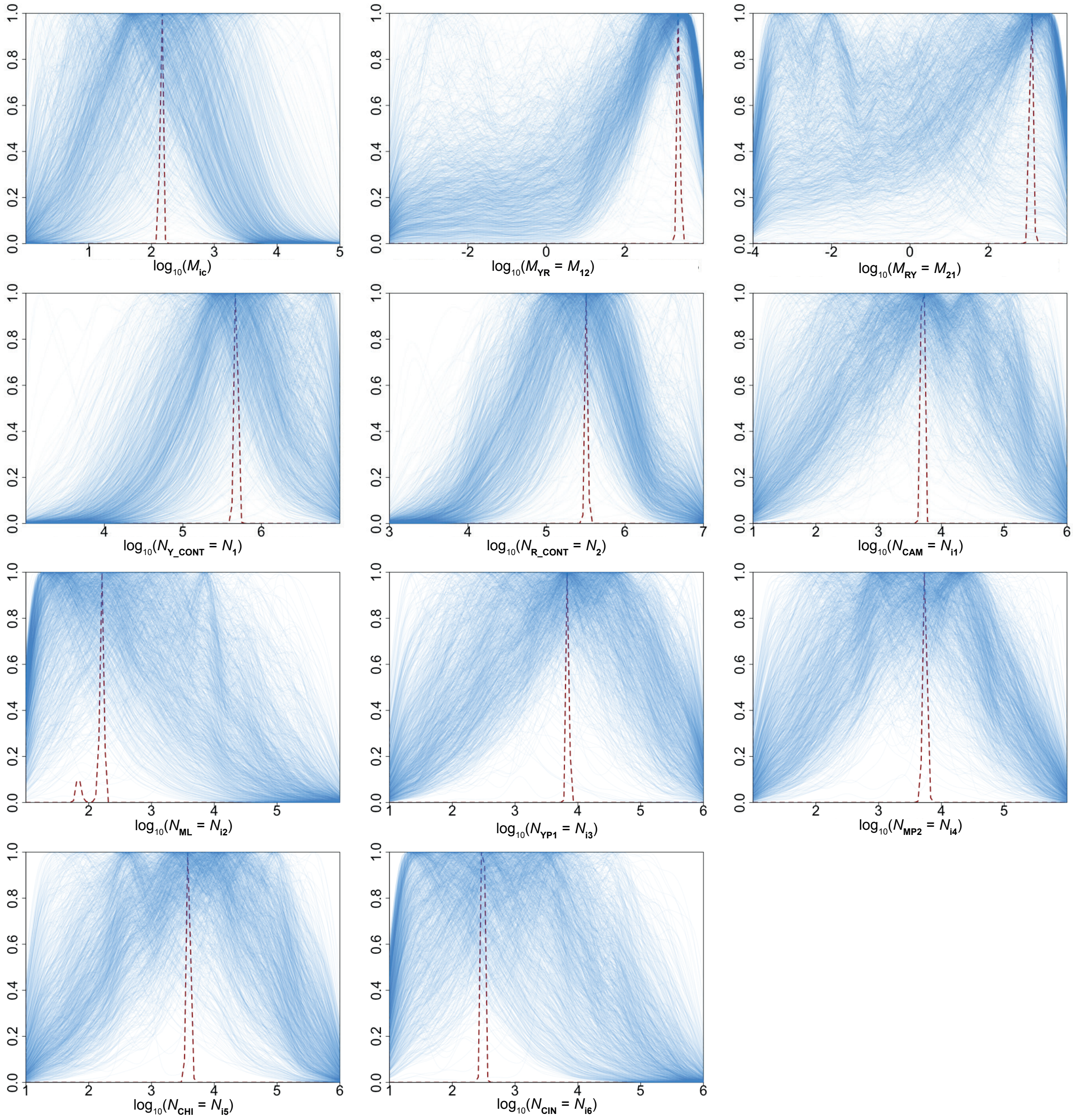


**Figure S16.** Estimation of $\widehat{\boldsymbol{\theta}}$ (red dashed lines) from the posteriors of 1191 10kb putatively neutral windows (blue solid lines) for model $\boldsymbol{\mathcal{M}_2}$ in the *A.majus* case study. Here, $M_{ic}$ are scaled to a global $N_E$ (=10,000), while $M_{12}, M_{21}$ are scaled to $\widehat{N}_1$ and $\widehat{N}_2$ respectively. $x$-axes ranges reflect prior ranges.
